# The Ingestive Response Reflects Neural Dynamics in Gustatory Cortex

**DOI:** 10.1101/2025.10.01.679845

**Authors:** Natasha Baas-Thomas, Abuzar Mahmood, Narendra Mukherjee, Kathleen C. Maigler, Yixi Wang, Donald B. Katz

**Affiliations:** Program in Neuroscience, Brandeis University, Waltham, Massachusetts 02454, USA; Department of Psychology, Brandeis University, Waltham, Massachusetts 02454, USA

**Author notes:** Corresponding author: Donald B. Katz. These authors contributed equally (Natasha Baas-Thomas, Abuzar Mahmood). **Author Contributions:** NBT, AM, and DK conceived and designed the research. NBT, NM, KM, and YW performed the experiments. NBT, AM, and DK interpreted the results of the experiments. NBT and AM prepared figures. NBT, AM and DK drafted and revised the manuscript. All authors approved the final version of the manuscript. **Competing interest statement:** The authors declare no competing financial interests.

## Abstract

Upon delivery of a taste onto the tongue, gustatory neural activity determines whether the stimulus is ingested or rejected. While some work in rodents has been devoted to investigating the neural activity leading to the rejection decision and its associated orofacial movements, little is known about what behaviors lead to ingestion of palatable tastes (and what neural activity is associated with that decision), largely because identifying ingestion-related behaviors is a difficult challenge—and probably undoable with video analysis given that the behaviors are largely intraoral. To address this gap in our understanding, we analyzed simultaneously-collected electromyographic (EMG) activity of the jaw opener muscle and the firing of gustatory cortical (GC) ensembles. We developed a machine-learning classifier to identify individual orofacial movements from EMG signals, demonstrating that it outperforms previously developed methods and using the technique to reveal three novel subtypes of ingestion-related tongue/mouth movements. Investigating the dynamics of these behaviors, we found that the frequency of occurrence of each type subtype shifts significantly at the time of the consumption decision, and is both tightly coupled with and reliably follows the transition in GC population activity into the state reflecting the tastant’s emotional/hedonic value. However, rather than the onset of single “ingestion” movement (as occurs for rejection decisions in the form of gapes), we show that the transition to ingestion is instead characterized by a collective change in the frequencies all ingestion-related behaviors. These findings demonstrate a direct link between neural dynamics in GC and the orchestration of the physical movements that define ingestive behavior, highlighting GC’s general role in taste perception, decision making, and the control of motor actions.

## Introduction

When a taste stimulus reaches the tongue, the animal has one basic goal—that of determining whether that stimulus should be ingested or rejected. In rats, consumption judgments are actuated by a family of orofacial behaviors generated by overlapping sets of lingual and masticatory muscles. A multi-functional central pattern generator in the brainstem drives the full range of these behaviors (Chen et al., 2001; DiNardo & Travers, 1997; Grill & Norgren, 1978b; Moriyama, 1987; Travers et al., 2000; Venugopal et al., 2007), but beyond this, much remains to be understood about the neural control of these consumption behaviors.

Of tastant rejection and consumption, the former has received far more study, partly because the orofacial behavior that defines the decision to expel a tastant is a large and stereotyped (hence, easily-detected and classified) triangular-shaped opening of the mouth called a “gape” (Grill & Norgren, 1978a). While gapes require the coordination of multiple muscles (as with ingestion), they can be reliably detected through electromyography (EMG) of a jaw opener muscle, the anterior digastric (AD), which contracts in direct correspondence to gape production. The ease with which gapes are detected has helped researchers relate rejection decisions to taste responses in the gustatory cortex (GC), which progress through three successive population activity “states,” the last of which contains firing correlated with tastant palatability (Jones et al., 2007; Moran & Katz, 2014; Sadacca et al., 2012, 2016). The transition into this palatability state reliably precedes the onset of gapes to an aversive taste by ~0.3 s (Li et al., 2016; Sadacca et al., 2016), and brief (0.5s) optogenetic inhibition of GC delays gaping if (and only if) the onset of the silencing perturbation occurs before the transition into the palatability state (Mukherjee et al., 2019), causally linking the GC transition to the initiation of this behavior. In summary, the GC ensemble transition into palatability-related firing participates in driving of rejection behaviors.

Several lines of evidence indirectly support the hypothesis that GC taste response dynamics also drive ingestive behaviors: first, GC dynamics in response to palatable and unpalatable tastants are distinct but similarly timed (Katz et al., 2001; Lin et al., 2021; Sadacca et al., 2012); second, perturbation of GC affects discrimination of palatable tastes in both rodents (Bales et al., 2015) and humans (Pritchard et al., 1999); third, GC projects both to feeding centers (such as amygdala and lateral hypothalamus, Saper, 1982) and to the above-mentioned brainstem region (Shammah-Lagnado et al., 1992; Zhang et al., 2011; Zhang & Sasamoto, 1990) that controls both ingestive and rejective orofacial movements (Berthoud & Münzberg, 2011; Chen et al., 2001; Moriyama, 1987; Travers et al., 2000).

This hypothesis has never been tested, however, because the set of motor behaviors comprising ingestion is far more complicated than gaping. In a seminal paper, Grill & Norgren (1978a) proposed that intraoral infusion of palatable tastants elicits a set of three distinct orofacial behaviors: lateral tongue movements (observed in response to only the most palatable tastants), tongue protrusions, and mouth movements. The latter two both involve rapid (6-8 Hz), low amplitude jaw opening (Grill & Norgren, 1978a; Travers & Norgren, 1986) with, in the case of tongue protrusions, extension of the tongue beyond the incisor plane (Grill & Norgren, 1978a). Although these behaviors resemble licking (the behavior elicited to acquire externally presented stimuli), they were proposed to capture stimulus-specific aspects of processing (Travers & Norgren, 1986).

However, there is a lack of consensus with regard to the classification and function of these behaviors: some experimenters have suggested alternative definitions of “mouth movements” as movements of the lower mandible without mouth opening (Jarrett et al., 2005; López et al., 2023; Parker et al., 1992), while others (Kaplan et al., 1995) have suggested that mouth movements and tongue protrusions, rather than being categorically separate behaviors, instead exist on a single continuum of tongue extension; still others have suggested that mouth movements are nearly (Berridge & Grill, 1983; Berridge & Treit, 1986; Kaplan et al., 1995) or completely hedonically neutral (Berridge, 1991; Brining et al., 1991; De Luca et al., 2012; Feurté et al., 2000), or not a part of the ingestive profile at all (Fontanini & Katz, 2006; Pereira et al., 2021; Soares et al., 2007). To avoid this ambiguity, some experimenters eschew the intricacies of ingestive behaviors altogether, focusing on how neural circuits drive “licking” (Chen et al., 2001; DiNardo & Travers, 1997; Travers & Jackson, 1992; Travers & Norgren, 1986).

Given this lack of clarity, testing the hypothesis that GC is involved in the driving of both ingestion and rejection requires a more sophisticated analysis of the EMG responses underlying ingestion. Here, we perform this analysis, using a machine learning classifier to systematically detect and categorize individual ingestive-related mouth and tongue movements (MTMs) in AD EMG activity. Our use of information-rich EMG data mitigates the difficulty of visually distinguishing mouth movements from tongue protrusions in unrestrained rats. This analysis reveals that MTMs are in fact made up of multiple novel behavioral subtypes that are at least partly distinct from those originally proposed by Grill & Norgren (1978a). Further analysis demonstrates that the relative frequency of each subtype changes suddenly within single trials, and that transitions in behavior align with neural transitions of palatability-related neural activity.

We conclude that consumption decisions are reflected in an ensemble of interacting behaviors, and that GC is involved not only in the generation of the rejection response, but also in behaviors related to ingestion. This further implicates GC in both processing gustatory stimuli and in effecting the subsequent motor response.

## Methods

### Experimental design and data collection

#### Subjects

Adult, female Long–Evans rats (*n* = 13, 250–400 g at time of surgical implantation, Charles River Laboratories) served as subjects in our study. A subset of these rats (n = 5) were taken from previously published data (Mukherjee et al., 2019; see **Table 1**). The rats were housed in individual cages in a temperature- and humidity-controlled environment under a 12 h light/dark cycle, given access to food and water *ad libitum* before the start of experimentation. All experimental methods were in compliance with National Institutes of Health guidelines and were approved in advance by the Brandeis University Institutional Animal Care and Use Committee.

**Table 1.** Summary of rat subjects and experimental conditions.

| Rat ID | # of Sessions | Tastants | Trials / Tastant | Data Types |  |  | Published |
| --- | --- | --- | --- | --- | --- | --- | --- |
|  |  |  |  | Anterior Digastric EMG | Video | # of GC Electrodes |  |
| NB27 | 3 | 0.3M Suc, 0.1M NaCl, 0.1M CA, 5mM QHCl | 30 | ✓ | ✓ | X | X |
| NB32 | 3 | Water, 0.3M Suc, 0.1M NaCl, 2mM QHCl | 30 | ✓ | ✓ | X | X |
| NB34 | 3 | Water, 0.3M Suc, 0.1M NaCl, 3mM QHCl | 30 | ✓ | ✓ | X | X |
| NB35 | 2 | Water, 0.3M Suc, 0.1M NaCl, 2mM QHCl | 30 | ✓ | ✓ | X | X |
| KM29 | 2 | 0.3 Suc, 0.1M NaCl, 0.1M CA, 1mM QHCl | 30 | ✓ | X | 14 | X |
| KM45 | 1 | Water, 0.3M Suc, 0.1M NaCl, 0.1M CA, 5mM QHCl | 25 | ✓ | X | 16 | X |
| KM50 | 2 | Water, 0.3M Suc, 0.1M NaCl, 0.1M CA, 5mM QHCl | 25 | ✓ | X | 16 | X |
| NB33 | 1 | Water, 0.3M Suc, 0.1M NaCl | 40 | ✓ | X | 30 | X |
| NB33 | 1 | Water, 0.3M Suc, 0.1M NaCl, 2mM QHCl | 30 | ✓ | X | 30 | X |
| NM51 | 1 | 0.3M Suc, 30mM Suc, 0.1mM QHCl, 1mM QHCl | 8 | ✓ | X | 30 | ✓ |
| NM51 | 1 | 0.3M Suc, 30mM Suc, 0.1mM QHCl, 1mM QHCl | 15 | ✓ | X | 30 | ✓ |
| NM50 | 2 | 0.3M Suc, 30mM Suc, 0.1mM QHCl, 1mM QHCl | 8 | ✓ | X | 30 | ✓ |
| NM50 | 2 | 0.3M Suc, 30mM Suc, 0.1mM QHCl, 1mM QHCl | 15 | ✓ | X | 30 | ✓ |
| NM47 | 1 | 0.3M Suc, 30mM Suc, 0.1mM QHCl, 1mM QHCl | 15 | ✓ | X | 30 | ✓ |
| NM47 | 2 | 0.3M Suc, 30mM Suc, 0.1mM QHCl, 1mM QHCl | 8 | ✓ | X | 30 | ✓ |
| NM44 | 1 | 0.3M Suc, 30mM Suc, 0.1mM QHCl, 1mM QHCl | 8 | ✓ | X | 30 | ✓ |
| NM44 | 1 | 0.3M Suc, 30mM Suc, 0.1mM QHCl, 1mM QHCl | 10 | ✓ | X | 30 | ✓ |
| NM43 | 2 | 0.3M Suc, 30mM Suc, 0.1mM QHCl, 1mM QHCl | 8 | ✓ | X | 30 | ✓ |
| NM43 | 1 | 0.3M Suc, 30mM Suc, 0.1mM QHCl, 1mM QHCl | 10 | ✓ | X | 30 | ✓ |
| NM43 | 1 | 0.3M Suc, 30mM Suc, 0.1mM QHCl, 1mM QHCl | 15 | ✓ | X | 30 | ✓ |

#### Electrode, EMG, and intraoral cannula construction

Custom microwire bundle drives were constructed to include between 0 (for EMG-only implants) to 30 recording electrodes (0.0015-inch formvar-coated nichrome wire; AM Systems; see **Table 1**). The microwire bundle was affixed to a custom-made electrode interface board (San Francisco Circuits) and connected to a 32-channel Omnetics connector. The remaining two pins of the connector were wired to electromyography (EMG) electrodes (PFA-coated stainless steel wire; AM Systems) for muscle implantation. Finally, the microwires were connected to a custom-built 3D printed microdrive that allowed the entire assembly to be moved ventrally after implantation. For more information on the implanted apparatuses and associated electronics, see (Katz et al., 2001; Li et al., 2016; Sadacca et al., 2016), as well as the Katz Lab webpage (https://katzlab.squarespace.com/technology).

Intraoral cannulae, flexible polyethylene tubing (AM Systems) with a flanged tip and washer to ensure stability, connected to a plastic top complete with a locking mechanism, were built to allow the delivery of tastants directly onto the tongue as described previously (Fontanini & Katz, 2006).

#### Surgical implantation

All subjects were implanted with an intraoral cannula (IOC) and EMG in the anterior digastric; some additionally had electrodes implanted in GC. All rats were first anesthetized with an intraperitoneal injection of a ketamine/xylazine mixture (100 mg/kg and 5 mg/kg body weight, respectively), after which we shaved and cleaned the scalp and under the jaw, and situated the head in the stereotaxic frame. After excising the scalp and leveling the skull, we drilled five self-tapping screws into the skull for supporting and grounding the electrode bundles.

For subjects that had an electrode implant, we next made an additional, larger craniotomy (∼2 mm dia.) above GC. An electrode microdrive (see *Electrode, EMG, and intraoral cannula construction*) was slowly lowered (over 5-10 min), and electrodes were positioned 0.5 mm above the GC (+1.3 mm A/P, +/-5 mm M/L, 4.6 mm D/V). Afterwards, the craniotomy was sealed with Kwik-Sil (World Precision Instruments).

For all subjects, the ground wires were wound tightly around the skull screws and the microdrive was cemented in place with dental acrylic. The rat was then removed from the stereotaxic frame and implanted with a single (right-side) IOC in the space between the first maxillary molar and the lip, through the masseter muscle and inside the zygomatic arch, and out through the opening in the scalp (Katz et al., 2001; Phillips & Norgren, 1970) before being cemented in place.

The EMG electrodes were channeled down the left side of the face (opposite from the IOC); after the overlying skin had been teased away from the belly of the digastric muscle, one end of each EMG electrode was then inserted into the muscle using a suture needle (for more details, see Dinardo & Travers, 1994; Li et al., 2016; Loeb & Gans, 1986; Travers & Norgren, 1986). The electrode wires were trimmed and held in place with vetbond tissue adhesive (3M) and the skin covering the anterior digastric was sutured closed. Finally, bacitracin ointment was applied all around the base of the headcap and over the incision under the jaw.

Rats were postoperatively injected with analgesic (buprenorphine 0.05 mg/kg or meloxicam 0.04 mg/kg), saline, and antibiotic (Pro-Pen-G 150,000 U/kg). Similar antibiotic, saline and analgesic injections were delivered 24, 48 and 72 hr later, and bacitracin ointment was reapplied. The rats were allowed to recover for a minimum of 6 days, and daily weight records were kept to ensure that the rats did not fall below 85% of pre-surgery weight.

#### Experimental design Habituation

Following recovery from the implantation surgery, we habituated rats to passive water deliveries in the experimental chamber for 2 to 3 days before beginning data collection. During these habituation sessions, 60 to 120 pulses of ~40 μL of distilled water were delivered into the animal’s oral cavity through the IOC. We also placed rats on a mild water restriction schedule, 15-20 mL of water (not including the 4 mL delivered during habituation sessions themselves) per day to ensure engagement in the task, which was maintained for the duration of the study. At the end of the habituation period, electrode bundles were driven deeper such that the electrode tips lay within GC.

#### Passive taste administration

Experimental test sessions spanned 1 to 4 days after habituation (1 session per day). Freely moving rats received three to five tastants *via* IOC in a pseudo-random order. Each tastant battery (see **Table 1**), selected from the following set, included a range of distinct taste identities and palatabilities comparable to those in our (Grossman et al., 2008; Levitan et al., 2019; Moran & Katz, 2014; Sadacca et al., 2012) and others’ (Samuelsen et al., 2012; Spector & Grill, 1988) studies: 0.1 M sodium chloride (NaCl), 0.3 M or 30 mM sucrose (Sucrose), 0.1 M citric acid (CA), 5 mM, 3 mM, 2 mM or 1 mM quinine-HCl (QHCl), 0.1 mM quinine-HCl (Dil-QHCl), or distilled water. Quinine-HCl concentrations (5-1 mM) and sucrose concentrations (0.3 M/30 mM) were each grouped, as these ranges elicit similar taste-responsive orofacial movements (Grill & Norgren, 1978a). During taste delivery, EMG and either video recordings or GC single unit activity were captured, as detailed below.

The taste delivery apparatus consisted of gently pressurized tubes containing taste solutions; the tubes converged upon a manifold of finer polyamide tubes that could be inserted into (and extending 0.5 mm past the end of) the IOC, thus eliminating any chance of mixing. The manifold could be locked securely into the dental acrylic cap. The tastes were then delivered under slight nitrogen pressure—this taste delivery protocol has been consistently shown to ensure reliable tongue coverage at short latencies (see Katz et al., 2001; Li et al., 2016; Sadacca et al., 2016).

#### Acquisition of electrophysiological and EMG data

Electrophysiological signals from the microelectrodes were sampled at 30 kHz using 32-channel analog-to-digital converter chips (catalog #RHD2132) from Intan Technologies, digitized online at the head stage and sampled jointly, along with signals from actuators marking tastant delivery, using an Intan RHD USB interface board (catalog #C3100), which routed records to the hard drive of a PC for saving. The experimental chamber was ensconced in a Faraday cage that shielded recordings from external electromagnetic influences.

#### Video recording

A camera (Brio 4K, Logitech) was positioned below the plexiglass experimental chamber to best record the orofacial movements of the subjects. The videos were captured with OBS Studio at a resolution of at least 1920 x 1080 and 60 frames per second. The delivery of a taste and a single red LED flash of 1 second were initiated simultaneously so that the light onset visually marked the start of the trial.

#### Histology

In preparation for histology to confirm GC electrode placement, rats were deeply anesthetized with an overdose of the ketamine/xylazine mixture. We perfused the rats through the heart with 0.9% saline followed by 10% formalin and harvested the brain. The brain tissue was incubated in a fixing mixture of 30% sucrose and 10% formalin for several days before being sectioned into 50 μm coronal slices on a sliding microtome (SM2010R, Leica Microsystems). Sections containing the electrode implant sites around GC were imaged with a fluorescence microscope (BX-Z710, Keyence). Histological verification for rats with only EMG implants was not necessary as targeting of EMG electrodes was directly observed during surgery.

### Data processing and statistical analysis

The analysis of data and statistical tests were performed using custom-written software in Python as described below. Some of the more prominently used packages are mentioned here:

- Scipy (statistical analysis; Virtanen et al., 2020)
- Pandas (data handling and manipulation; The pandas development team, 2025)
- Scikit-Learn (machine learning; Pedregosa et al., 2011)
- Numpy (numerical computing; Harris et al., 2020)
- Pingouin (hypothesis testing; Vallat, 2018)
- Blech_clust (electrophysiological data processing; Mahmood et al., 2025)
- Scikit-Optimize (hyperparameter optimization; Head et al., 2024)
- UMAP (non-linear dimensionality reduction; McInnes et al., 2018)

#### Video scoring

##### Scoring methodology and ethogram

Videos of each session were first reviewed to identify instances in which the rat’s mouth was clearly visible (i.e., not obstructed because of odd head angles or forepaws resting in front of the mouth) following taste delivery. The selected trials were then analyzed frame-by-frame to classify and record the subject’s orofacial movements using BORIS (Friard & Gamba, 2016). All scoring was performed blind to the delivered taste to minimize potential bias. Two scorers independently evaluated the presence of three taste-responsive orofacial movements: gape (retraction of the corners of the mouth, resulting in a wide triangle-shaped opening of the mouth), mouth and togue movements (MTMs, rhythmic, low-amplitude openings of the mandible, with or without tongue emergence), and lateral tongue movement (lateral extension of the tongue). Note that mouth movements (rhythmic, low-amplitude openings of the mandible) and tongue protrusion (emergence of the tip of the tongue directly on the midline during mouth opening) are combined into one behavior: MTM, because in video recordings it was not possible to reliably distinguish whether or to what extent the tongue emerged, making it difficult to separate these behaviors. Difficulty in distinguishing these behaviors was further exacerbated by the lack of consensus in the literature regarding the discriminative characteristics of these two behaviors (see Introduction). Periods of no movement (absence of any perceptible orofacial muscle activity) were also marked by the scorers, though this category was not assessed for inter-rater accuracy due to its simplicity.

Since video sections were focused on taste responsive sections, we supplemented the number of “no movement” samples by using the EMG signal to select pre-stimulus (1000-2000 ms pre-stimulus) activity trials with no discernible orofacial movements (based on the amplitude of the pre-stimulus signal relative to the average post-stimulus signal amplitude) and incorporating all segmented behaviors from that time period as “no movement” samples.

##### Interrater scoring accuracy

To assess inter-rater accuracy, we analyzed the first 5 seconds of trials that were independently scored by both observers. In each comparison, one scorer’s behavior annotations were treated as the “ground-truth,” and we evaluated whether the second scorer had a temporally overlapping annotation of the same behavior type. Any overlap in annotation was considered an agreement; annotations without any overlap were considered disagreements. Accuracy was computed separately for each behavior category (MTMs, gapes, LTMs). This analysis was then repeated with the second scorer treated as the ground truth. Agreement accuracy was then calculated as the proportion of matched bouts out of the total number of bouts for each scorer. This can be formalized as:

Accuracy 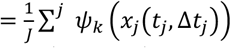

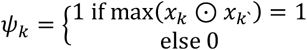

*i* = Counter over label categories

*j* = Counter over label instances for a given labeler

*k* = Counter over labellers

*x*(*t* _*j*_,Δ*t* _*j*_) is a binary vector representing the duration of a specific labelled event

To estimate a chance level of agreement, we repeated the analysis 100 times with randomized annotations. Specifically, in each shuffle iteration, the behavior annotations of a scorer were randomly reassigned new labels (uniformly sampled between MTM, gape, and LTM), after which, agreement accuracy was recalculated.

#### Electrophysiology

##### Single unit isolation

Spikes from electrophysiological recordings were sorted and analyzed off-line using an in-house Python-based pipeline (Mahmood et al., 2025). Putative single-neuron waveforms with >5:1 signal-to-noise ratio (median absolute deviation) were sorted using a semi-supervised algorithm: recorded voltage data were filtered between 300-3000 Hz, grouped into potential clusters by a Gaussian Mixture Models (GMM) fit to multiple waveform features; clusters were then labeled and/or refined manually (to increase conservatism) by the experimenters.

##### Changepoint Estimation

We used an independent multivariate Poisson changepoint model implemented in the pytau package (Mahmood, 2026) to infer when changes in the probability of spike-counts occured in the timeseries of taste-evoked neural responses as done previously (Flores et al., 2025; Mahmood et al., 2023, 2026).

Models were fit using Automatic Differentiation Variational Inference (ADVI) functionality in the pymc library for 1e5 iterations (Salvatier et al., 2016). Briefly, the structure of the model is given below. The subscripts following the distribution denote the size of the variable.

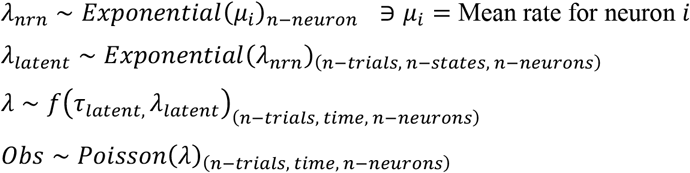

f = A function which combines tau-latent and lambda-latent to generate the inferred probability for population activity for each trial and timestep.

##### Determining best number of states for neural models

We fit changepoint models with 2-8 states to all datasets using ADVI as described above. ADVI optimizes the Evidence Lower Bound (ELBO), which is an estimate of the marginal likelihood of the model, providing an estimate of goodness of fit while accounting for model complexity (number of parameters), in a similar way to frequently used information criteria as AIC and BIC (Akaike, 1998; Dziak et al., 2019). Lower ELBO values indicate better model fitting.

#### Electromyography

##### EMG processing

As detailed previously, we recorded voltage signals from two unipolar EMG electrodes implanted in the anterior digastric muscle at 30 kHz. We used the difference in the voltage recorded by the two electrodes as the EMG signal—this procedure helps to cancel out any large artifacts produced by the animal’s movements and is equivalent to using a differential amplifier (as done in Li et al., 2016). We down-sampled the EMG signal to 1000 Hz by averaging the voltage values in sets of 30 samples, and high pass filtered the down-sampled signal above 300 Hz (Li et al., 2016; J. B. Travers & Norgren, 1986) using a 2^nd^ order Butterworth filter. The absolute value/magnitude of the filtered EMG signal was then lowpass filtered (again using a Butterworth filter of order 2) below 15 Hz, effectively capturing the envelope of variation of the EMG signal (plotted as the black curve in **Figure 1A**). This cutoff of 15 Hz is sufficient for identifying orofacial behaviors, all of which occur at frequencies smaller than 10 Hz (Li et al., 2016; J. B. Travers & Norgren, 1986).

**Figure 1.**
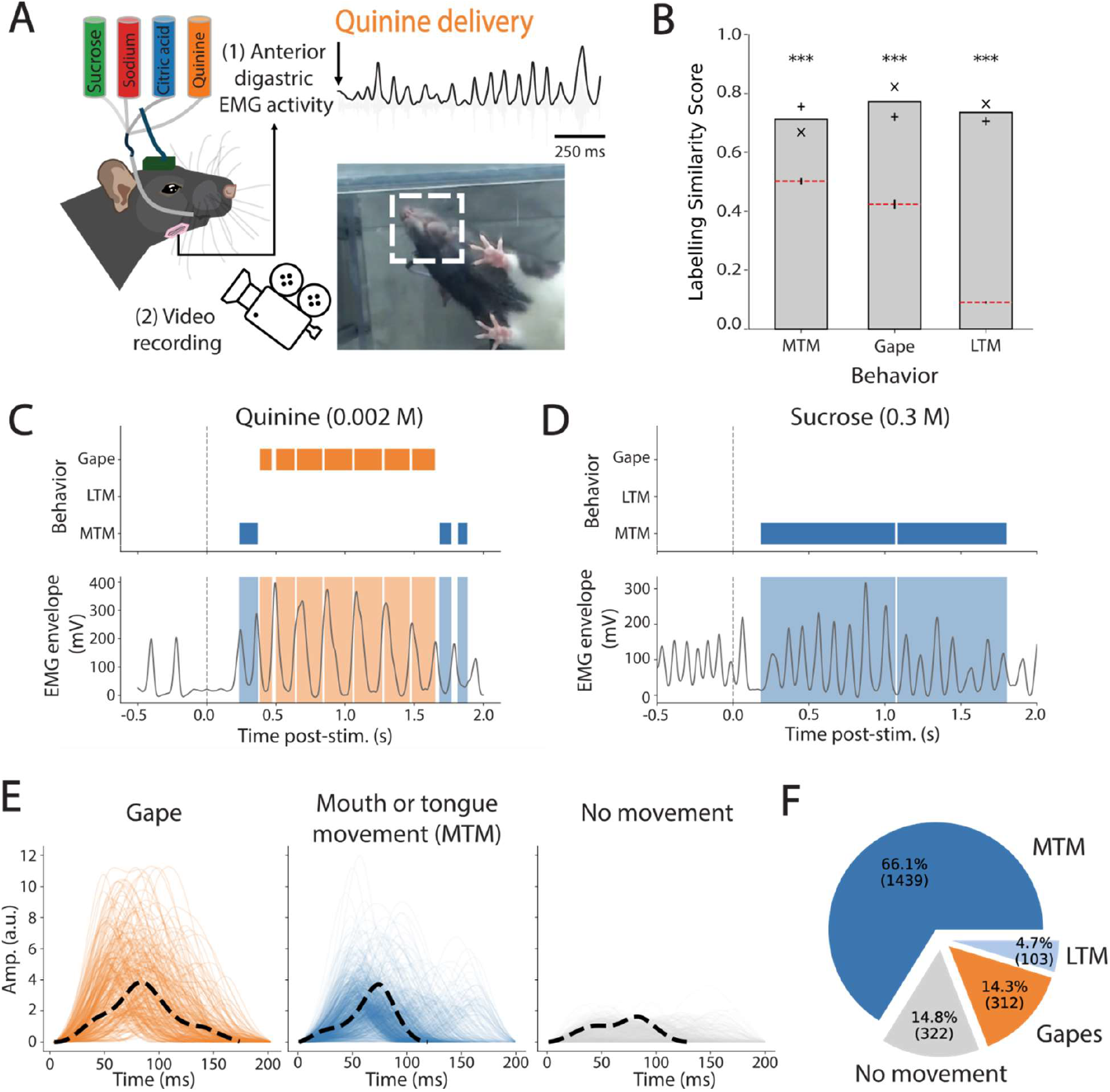
Annotation of EMG waveforms using scored video labels. **A)** A schematic of the data recording and annotation setup. Rats implanted with electromyography (EMG) electrodes and an intraoral cannula were given a battery of tastants while EMG and video recording were captured.**B)** Video scoring similarity between the two raters was significantly higher than chance for all labels (*** p < 0.001, one-tailed t-test). x, + = accuracy score with scorer 1,2 as ground-truth respectively. **C-D**) Examples of video-scored behaviors (top), and corresponding EMG signals (bottom) for trials of aversive quinine (C) and palatable sucrose (D). Note different y-scales for EMG envelopes between C and D. **E)** Overlain waveforms for each group of labelled movements (Lateral Tongue Movements [LTMs], which made up <5% of behaviors, are not shown). **F)** Breakdown of the labelled dataset by behavior.

##### XGB Classifier

Code for generating and running the classifier, as well as the final training dataset can be found at https://katzlabbrandeis/blech_emg_classifier

##### Movement extraction and refinement

We used low-pass filtered EMG signals (as described above) to extract individual orofacial movements by first applying a top-hat filter (scipy, length = 200 ms) to the signal and segmenting contiguous non-zero regions of the filtered signal. In some cases, this resulted in waveforms which had biologically implausible durations (too long or short). We removed any extracted waveforms shorter than 50 ms and longer than 500 ms.

##### Removal of multimodal MTM waveforms

We found that our process extracted some MTMs waveforms that were multimodal (3.9%), likely due to two muscle contractions that were improperly segmented. Despite their small fraction, these waveforms added substantial variability to the data and therefore needed to be removed. To assess multimodality, we performed a positive-lagged autocorrelation. The presence of one or more peaks (scipy.signal.find_peaks) indicated that the waveform contained repeating structure consistent with multimodality. Waveforms identified as multimodal were removed from the dataset and from all subsequent analyses.

##### Expansion of no-movement category

Due to the noisy nature of EMG activity, as well as the imperfect labelling and extraction of individual orofacial movements, we observed taste-related orofacial movements (mostly MTMs but also some gapes) with very small amplitudes, likely due to the unavoidable labelling of all orofacial movements within a range of time from video scoring. To address this issue, we absorbed /relabeled mouth movements most similar to the “no movement” category in feature space. To do this, we performed Neighborhood Component Analysis (NCA) to extract the 2 dimensions which following a linear transformation best allow classification of all 3 mouth movement categories using a Nearest Neighbor Classifier. In the 2D-NCA space, we fit Gaussian distribution to the “no movement” category and reclassified all mouth movements which were within 2 Mahalanobis distances of this Gaussian.

##### Feature engineering

We extracted a total of 8 features for each individual mouth movement. Two of these— duration and time to next peak—were based on the previously published QDA classifier (Li et al., 2016), and another—maximum frequency—was based on the previously published BSA classifier (Mukherjee et al., 2019). We included 5 additional features—amplitude, time to previous peak, and the first 3 Principal Components of the amplitude and duration normalized waveforms (to extract waveform shape-related features)—to assess whether these improved classification performance. PCA was trained using waveforms from the entire training dataset. All features were then standardized using the StandardScaler implemented in Scipy. Such extraction of features is standard practice to improve computational efficiency and model fitting, especially for clustering algorithms by reducing the dimensions representing each waveform (Khalid et al., 2014). Reducing the dimensionality of the data combinatorially reduces computational complexity and decreases the sparsity of the representational space of the data, allowing clustering algorithms to more effectively discover spatial patterns (Radovanović et al., 2010). Both the fitted PCA and StandardScaler objects were then frozen (dumped) and are available for reuse from the GitHub repository.

##### Hyperparameter optimization

To ensure robustness, we tested that the training of our classifier was agnostic to hyperparameters specification. We used the scikit-optimize library (Head et al., 2024) to perform 20 optimization runs with 400 iterations/run using leave-one-animal-out cross-validation accuracy as the optimization target, and the GBRT (Gradient Boosting Regression Trees) model as a surrogate function to sequentially model the fitness landscape (**Table 2**). The chosen set of hyperparameters is available in the GitHub repository.

**Table 2.** Optimization space of classifier hyperparameters.

| Name | Domain | Distribution | Min | Max |
| --- | --- | --- | --- | --- |
| learning_rate | Real | log-uniform | 0.001 | 1.0 |
| min_child_weight | Integer | Discrete uniform | 1 | 10 |
| max_depth | Integer | Discrete uniform | 1 | 100 |
| max_delta_step | Integer | Discrete uniform | 0 | 20 |
| subsample | Real | Uniform | 0.01 | 1.0 |
| colsample_bytree | Real | Uniform | 0.01 | 1.0 |
| colsample_bylevel | Real | Uniform | 0.01 | 1.0 |
| reg_lambda | Real | log-uniform | 1e-9 | 1000 |
| reg_alpha | Real | log-uniform | 1e-9 | 1.0 |
| gamma | Real | log-uniform | 1e-9 | 0.5 |

##### Determining important features

We assessed the importance of features using SHAP analysis (Lundberg et al., 2020). Briefly, SHAP (SHapley Additive exPlanations) is a game theoretic approach to explain the output of any machine learning model. It connects optimal credit allocation with local explanations using the classic Shapley values from game theory and their related extensions. We used the Mean Absolute SHAP values implemented in the SHAP package (https://github.com/shap/shap) as a metric for feature importance.

##### Evaluation Classifier Performance

In order to quantitatively evaluate the performance of our classification model, we employed the standard metric of Accuracy, along with three additional metrics: precision, recall, and F1-score to obtain a detailed comparison of the models.

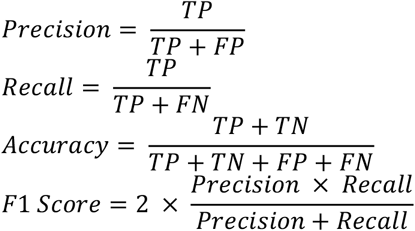

Where TP=True Positive, FP = False Positive, FN = False Negative, FP = False Positive Precision (also known as Positive Predictive Value) assesses the model’s accuracy among its positive predictions, effectively answering the question “Of all the events the model identified as positive, how many were actually positive?”. A high precision value indicates a low rate of false positives, which is critical in applications where such errors are costly.

Recall (also known as Sensitivity) measures the model’s ability to find all the positive events within the data, addressing the question “Of all the actual positive events in the data, how many did the model correctly identify?”. A high recall value indicates a low rate of false negatives, which is particularly important in screening applications where missing a positive case is undesirable. Given the inherent trade-off between precision and recall, a model optimized for high precision may miss many true positives, while a model optimized for high recall may generate many false positives.

To provide a single, balanced metric that accounts for this trade-off, we calculated the F1-score. The F1-score is the harmonic mean of precision and recall, providing a more robust measure of overall model performance, especially in datasets with an imbalanced class distribution. This score ranges from 0 to 1, with a value of 1 representing perfect precision and recall. The F1-score is therefore a suitable metric for evaluating a model where both false positives and false negatives are of significant concern.

We note precision, recall, and F1 score overlap with the commonly used d’ (Sensitivity Index or Fisher Discriminant Index) which is an unbiased estimator of the capacity of a classifier for binary classification calculated according to the following equation.

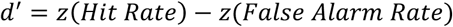

Where Hit Rate is the same as the True Positive Rate, and the False Alarm Rate is the same as the False Positive Rate.

##### Specificity to post-stimulus period

To assess consistency of model predictions to the taste-responsive period (previously shown to be primarily between 0-2000 ms post-stimulus delivery, Katz et al., 2001; Sadacca et al., 2012), we calculate for gapes and MTMs the fraction of events observed between 0-2000 ms post-stimulus delivery out of the total prediction window of -1000-3000 ms post-stimulus delivery. Logically, these taste-specific behavioral responses should only occur following stimulus delivery and high occurrence prior to stimulus, as labelled by a specific classifier, indicates a higher rate of false positives.

### Analysis of mouth or tongue movements (MTMs)

#### Clustering MTMs into behavioral subtypes

Events classified as MTMs were analyzed on a session-by-session basis. Principal components retaining 90% of the variance of the scaled MTM features were extracted; while preserving all original dimensions did not affect the final optimal number of clusters (data not shown), dimensionality reduction was retained to improve model fitting quality and computational efficiency. These reduced features were then clustered using Gaussian Mixture Model (GMM) with model selection performed using the Bayesian Information Criterion (BIC) to select the optimal number of clusters between 1 to 14. The optimal BIC score was determined by finding the “elbow”, i.e. the single breakpoint of a piecewise linear regression. This process was repeated for 50 iterations for each session to mitigate chances of being stuck in a local minimum.

This procedure revealed the mean optimal cluster number to be 3; hence we performed downstream analyses with 3-cluster models. We verified that the clusters were non-overlapping (as is allowed by GMMs) by calculating the pairwise Mahalanobis distances between all clusters, and confirmed that inter-cluster distances were significantly larger than intra-cluster distances using a Kolmogorov-Smirnov test.

#### Alignment of MTMs cluster labels across sessions

We aligned the labels across test sessions because cluster index labels are randomly assigned during the clustering process above. We performed this alignment using a “greedy” (sequential) approach. We first calculated the average feature vector for each cluster from two sessions and computed the cosine similarity between each pair. Cluster labels were then reordered to maximize the summed cosine similarity. The three clusters were then merged across sessions, and the resulting clusters were compared to the next session. This process was repeated, comparing the merged set of clusters to a new session, until all sessions were aligned.

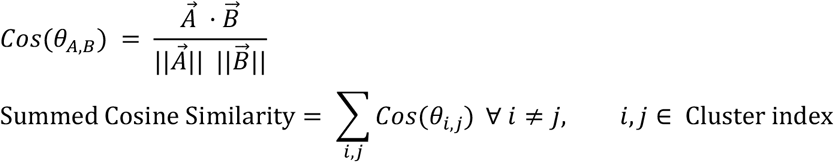

#### Support vector machine verification of MTM subtype labels

As further validation that the MTM clusters were separable and reliable across animals, we trained a support vector machine with a radial basis function kernel (scikit-learn.svm) on the 8 waveform features (see “Feature engineering”, Methods) using a leave-one-session-out cross-validation scheme. For each iteration, one session was designated as the test set, and all data from the corresponding animal were excluded from the training set. Predictions were generated on the held-out session, and classification accuracy was computed. To assess whether accuracy exceeded chance (33%), a one-tailed t-test was performed.

#### Analysis of MTM waveform feature differences across clusters

We extracted seven additional features from each non-normalized waveform segment to further characterize how waveform shapes differed between clusters: amplitude, area under the curve, full width half maximum, symmetry, skew, rising slope, and falling slope. Of note, symmetry was quantified as the Pearson correlation between the waveform and its mirrored version, and skewness was calculated using the sample skew statistic (scipy.stats.skew). Feature differences across clusters were evaluated with the Kruskal-Wallis test, and significant features with substantial effect sizes were followed up with a Mann-Whitney U-Test.

#### Temporal analysis of behavioral event counts

The three MTM subtypes between 0.3 s to 1.3 s post-stimulus delivery (a 1 s window around the average time to palatability-related firing in GC) were isolated for every trial. Behaviors were counted as either “before” or “after” the 800 ms mark. We assessed the differences in the distribution of MTMs observed “before” vs “after” by first projecting the waveforms from each session into 2D-space using UMAP, generating “before” and “after” distributions by binning them into histograms, subtracting the “before” from “after” binned density histograms, summing the absolute values of non-zero bins to use as our difference statistic. A null distribution for this difference statistic was generating using one thousand bootstrap shuffles of the MTM events (shuffling “before” and “after” labels) and p-values were calculated relative to this null distribution. As UMAP is a stochastic embedding (i.e., repeats of the procedure need not look the same), we performed this procedure 10 times (obtaining 10 p-values) to ensure robustness of our results.

For a more fine-grained comparison of gapes and three MTM subtypes, for each test session, a Poisson means test was used to determine whether movement counts differed significantly between the “before” and “after” groups for every taste. To assess whether the number of significant changes collectively exceeded the proportion expected by chance with an alpha of 0.05 (5%), we generated a null distribution of p-values using a shuffling procedure. A binomial test was used to test whether the number of significant p-values observed was significantly greater than expected by chance.

#### Changepoint estimation

We used a Categorical changepoint model implemented in the pytau package (Mahmood, 2026) to infer when changes in the probability of occurrence happened in the timeseries of clustered oral behaviors. Model fitting and comparison were performed in the same manner as for neural data (see Electrophysiology). The structure of the model is briefly described as follows:

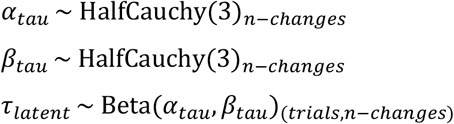

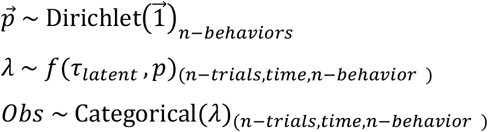

n-changes = n-states - 1 (both are used for clarity, above) f = A function which combines tau-latent and p to generate the inferred probability for each behavior for each trial and timestep.

#### Estimating the contribution of gapes and no movements to state transitions

We reasoned that if gapes and no movements were driving state changes, then removing them would significantly increase the similarity of subsequent states, making state transitions more difficult to infer. To perform this calculation, we used the mean occurrence probabilities of behaviors in each state inferred by the changepoint model, defined as follows:

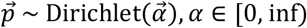

As a measure of dissimilarity, we calculated the L_2_ norm of the difference in probability vectors between subsequent states.

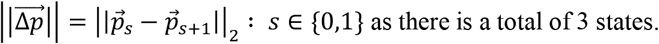

We standardized the distribution of these 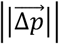 for all behaviors and for only MTMs using the mean norm difference expected by random vectors of corresponding sizes to account for the difference in the dimensionality of the vectors.

### Analysis of neural and behavioral changepoint alignment

#### Data Filtering

As shown previously, inference of sharp changes (transitions) in neural responses is sensitive to 1) the # of neurons in the dataset (Jones et al., 2007), 2) the number of neurons that participate in a given transition (Jones et al., 2007), and 3) the stability of the neural response across repeated stimulus presentations (Svedberg & Katz, 2024). To minimize the impact of these factors on our analysis, we performed changepoint analysis only on units that passed the following criteria:

1. Stability during an experimental session, tested using a 2-way ANOVA with factors time-bins (4 x 500ms bins of post-stimulus activity) and trial-bins (trials in a session divided into 4 bins) with the requirement that there was no main effect for trial-bins (p>0.05).
2. Either of:
  a. Stimulus responsiveness (p<0.05 for paired t-test on average firing for 1000ms pre-stimulus and 1000ms post-stimulus)
  b. Stimulus specificity or dynamicity in the stimulus-evoked response (p<0.05 for either factor in a 2-way ANOVA with factors = [time-bins, stimulus-identity], where time-bins:= [2000ms post-stimulus activity divided into 4 bins])

We implemented an additional model-level filter where we required that for each session, a model-comparison for models with 2-8 states (see section “Determining best number of states for neural models”)showed that models with 3 or 4 states fit the data best in-line with previous work (Jones et al., 2007; Mahmood et al., 2023, 2026), indicating both good inference of these well-established dynamics as well as minimal drift in neural responses which necessitates the use of additional states by the model. Finally, we only used trials with no optogenetic perturbations.

We note that all above-mentioned filtering is orthogonal to the desired analysis, that is, to test the alignment of neural and behavioral changes. The existence of these dynamics and transitions in neural responses is thoroughly established (Escola et al., n.d.; Jones et al., 2007; Mahmood et al., 2023, 2026; Moran & Katz, 2014; Mukherjee et al., 2019; Sadacca et al., 2016; Svedberg & Katz, 2024) and is not being tested here. The implemented filters instead aim to mitigate the confound of poor model-fits leading to a false-negative outcome.

#### Hypothesis testing for alignment

Using inferred changepoints for ephys and behavioral data, we performed a Pearson’s R correlation to test whether the latency of matching changepoints of each datatype were related (that is, Neural changepoint 1 was compared to behavior changepoint 1, and same for changepoint 2). An alpha=0.05 was used to indicate a significant relationship. To aggregate comparisons across the entire dataset, we tested whether the number of significant correlation was higher than that expected by chance using a binomial test (alpha=0.05), similar to previous work (Mahmood et al., 2023).

#### Changepoint Lag Analysis

In line with results from previous work showing that neural responses precede behavioral changes (Sadacca et al., 2016), we tested the consistency of lags between neural dynamics (i.e., coherent ensemble firing-rate transitions) and behavioral changes in our data. We used 3 sets of tests to ensure our results were not dependent on the “scale” of our statistics. First, we performed paired t-tests on changepoints pooled across all considered sessions; that is, for changepoint 1, we compared neural and behavioral changepoints from trials pooled across sessions. We followed this by performing a paired t-test on session-averaged latencies, calculating for each session the average latency for neural and behavioral changepoints and comparing them against each other. Finally, we assessed paired t-tests on a single session basis; that is, for every session we performed a paired t-test comparing neural and behavioral changepoint latencies for each changepoint index. We assessed the aggregate significance of these multiple paired t-tests by employing a Binomial test to assess whether the count of significant test was higher than that expected by chance.

#### Calculating state-specific palatability correlation

In line with results from previous work showing that the emergence of palatability-related activity in GC occurs in a state-specific manner (Sadacca et al., 2016), we tested whether the same is true in our data. We used state-durations inferred as part of the above analyses to calculate the average firing of neurons for any given state across all presentations of stimuli (tastants). The canonical palatability rank for each tastant was determined using the volume of each tastant consumed in a Brief Access Task. These ranks were correlated to the state-averaged neural activity of each neuron to determine the palatability “content” of each state. As a control, the same correlation was performed after shuffling trial-labels.

## Data availability

We have structured our electrophysiology datasets in a hierarchical data format (HDF5) and are hosting the files on a university-wide network share managed by Library and Technology Services (LTS) at Brandeis University. These HDF5 files contain our electrophysiology recordings, sorted spikes, and single-neuron and population-level analyses (and associated plots and results). These files are prohibitively large to be hosted on a general-purpose file share platform; we request people who are interested in our datasets to contact the corresponding author, Donald Katz, who can put them in touch with LTS to create a guest account at Brandeis University to securely access our datasets (hosted on the katz-lab share at files.brandeis.edu). As noted above, training data for the classifier is available as part of the GitHub repository.

## Results

### Overview

Our investigation into whether ingestive-related orofacial movements change in relation to GC dynamics contained 3 main parts.

First, we developed a machine-learning classifier that enabled us to identify individual orofacial movements, all of which cause some degree of jaw movement, from AD EMG activity. We began by labeling a subset of EMG waveforms, each representative of a single AD contraction, as either gapes, MTMs, or no movement using blind-scored videos. A gradient-boosted, tree-based classifier was then trained on features extracted from these labeled waveforms. We validated the classifier’s accuracy, ensuring it performs well on out-of-sample (novel) data, and confirmed that its performance matched or exceeded previously published gape detection algorithms (Li et al., 2016; Mukherjee et al., 2019).

Second, having verified that we could reliably classify both ingestive and aversive orofacial movements, we conducted a detailed analysis of MTMs in responses to palatable tastes using labels inferred by the classifier, which revealed MTMs to be comprised of three distinct and previously undescribed behavioral subtypes. We resolved subtle differences in the EMG properties of these MTM subtypes that would be difficult to distinguish from video, and observed changes, across the course of a single trial, of the emission frequency of all three subtypes.

Finally, we tested whether these within-trial changes in emission of MTM subtypes align with decision-related GC dynamics. Evaluation of GC neural ensembles recorded simultaneously with AD EMG activity revealed that changes in MTM subtype frequencies follow the onset of palatability-related firing in GC at short latencies (despite large trial-to-trial variability in both). This result provides evidence that GC is involved in initiation of both ingestive and rejective behaviors.

### Video scoring provides a training dataset of individual AD EMG waveforms

In order to reliably assess ingestion-related mouth movements, we developed a machine learning classifier capable of discriminating ingestive and rejective behaviors using AD (jaw opener) EMG activity. We generated a training dataset for the classifier by applying labels of taste-responsive behaviors identified from videos by expert scorers to AD-EMG waveforms (see **Fig. 1A** for experimental setup).

This training dataset was comprised of nine sessions in which freely moving rats (n = 4) were presented with a battery of basic tastants (0.3 M Sucrose, 5 mM Quinine, 0.1 M Citric Acid, and 0.1 M Sodium Chloride) *via* an intraoral cannula. Video recordings (sampling rate = 60 Hz) and AD EMG activity simultaneously captured orofacial responses to the tastes (**Fig. 1A**). Two video scorers, both blind to tastant identity, independently evaluated the subset of trials in which the rat’s mouth was clearly visible (13% of the total number of trials). The scorers labeled time points (within 5.1±3.7 s of post-stimulus activity, mean±SD) at which consummatory-related behaviors could be identified. The ethogram used for scoring followed mimetic descriptions established by Grill and Norgren (1978a), with one modification: to enhance inter-rater reliability (see Methods), mouth movements and tongue protrusions were combined into a single behavioral category, “mouth or tongue movements” (MTMs), defined as rhythmic, low-amplitude openings of the mandible, with or without tongue emergence.

Consistency of video labels from the two scorers was assessed using an accuracy classification score which accounts for the temporal component of video labels (a component that standard inter-rater reliability metrics typically neglect). This analysis revealed that inter-scorer similarity of labels was significantly higher than expected by chance (p < 0.001 for all comparisons, one-tailed t-test) for all three behaviors (MTMs: accuracy = 0.712, shuffle = 0.50±0.012; gapes: accuracy = 0.772, shuffled = 0.42±0.016; LTMs: accuracy = 0.735, shuffle = 0.09±0.005; mean±SD, **Fig. 1B**), demonstrating that the scorers were reliably classifying behaviors. The quality of our labelling is further corroborated by the fact that our labelled gape waveforms show dominant frequencies which match excellently with classification bounds used by previous studies (**Supplementary Fig. 1**, see Mukherjee et al. 2019).

Time-synced EMG records of AD activity were then aligned to the video record, such that time segments corresponding to the user-scored behavior labels could be matched to waveforms extracted from the EMG signal. **Figures 1C** and **1D** present examples of this analysis—behavioral events occurring within 2 seconds of taste deliveries along with the corresponding EMG activity. Peaks in EMG activity (previously identified as driving single orofacial movements, Li et al., 2016; Mukherjee et al., 2019) were identified and used to extract waveforms (**Figure 1E**). MTMs comprised a large majority of these behaviors (66.1% of a total n = 2,176) while gapes constituted 14.3%. We also isolated an equivalent number of “no movement” waveforms (14.8%; **Figure 1F**). As there were expectedly very few identified LTMs (4.7%)—too few for us to train as a distinct category for classifier training—these were excluded from further analysis.

### A machine-learning classifier accurately captures each class of orofacial movements in EMG signals

To infer behavioral responses from EMG traces, previous studies have either performed visual analysis (Dinardo & Travers, 1994; Sasamoto et al., 2001; J. B. Travers & Norgren, 1986), developed classifiers to assign labels to individual orofacial movements (Quadratic Discriminant Analysis [QDA], Li et al. 2016), or identified the probability of a rhythmic train of events using solely the frequency of the signal (Bayesian Spectrum Analysis [BSA], Mukherjee et al. 2019). These classification algorithms share the assumption that an orofacial movement can be indexed and discriminatively identified by an AD contraction: while any movement recruits multiple muscles, each behavior involves some movement of the jaw, and these movements are distinctive. As such, we refer to each AD waveform as representing a behavior.

The QDA and BSA classifiers were trained and validated purely for the prediction of gapes (Li et al., 2016; Mukherjee et al., 2019), and they performed well because gaping is an easily isolated and already well-understood behavior. However, for this same reason, they are unlikely to perform well (or be at all suitable) with other, more subtle behaviors (as we demonstrate below). To address these limitations, we developed a more complex classification model based on a gradient-boosted trees algorithm (XGB, Chen & Guestrin, 2016) capable of performing classification of the extended set of the labelled EMG signals of orofacial movements.

Rather than directly training the classifier on the EMG waveforms, we first extracted a suite of characteristics (features) from each waveform, a step that improved the computational efficiency of the classifier and allowed us the opportunity to directly interpret differences in features between differently classified orofacial movements (in line with previous work, see Li et al., 2016; Mukherjee et al., 2019; see also Methods for a discussion on how such “feature engineering” is a standard way to address dimensionality issues).

In applying this method, we make use, in a unified and principled framework, of the range of information sources that have been used in previous forms of EMG analysis. The use of a high-capacity tree-based model provides excellent interpretability (see **Fig. 2A**, a schematized version of the algorithm as a mixture of trees) and the opportunity to explore whether incorporating previously ignored features improves classification accuracy. We extracted the following features: amplitude; multiple frequency-based properties including maximum frequency, time from previous peak, time to following peak, and duration; and shape-related properties extracted using the first three principal components of standardized waveforms from the training set. We then applied SHAP-based model interrogation (<u>SH</u>apley <u>A</u>dditive ex<u>P</u>lanations, Lundberg et al. 2020) to test the contribution of each feature to the predictions made by the model.

**Figure 2.**
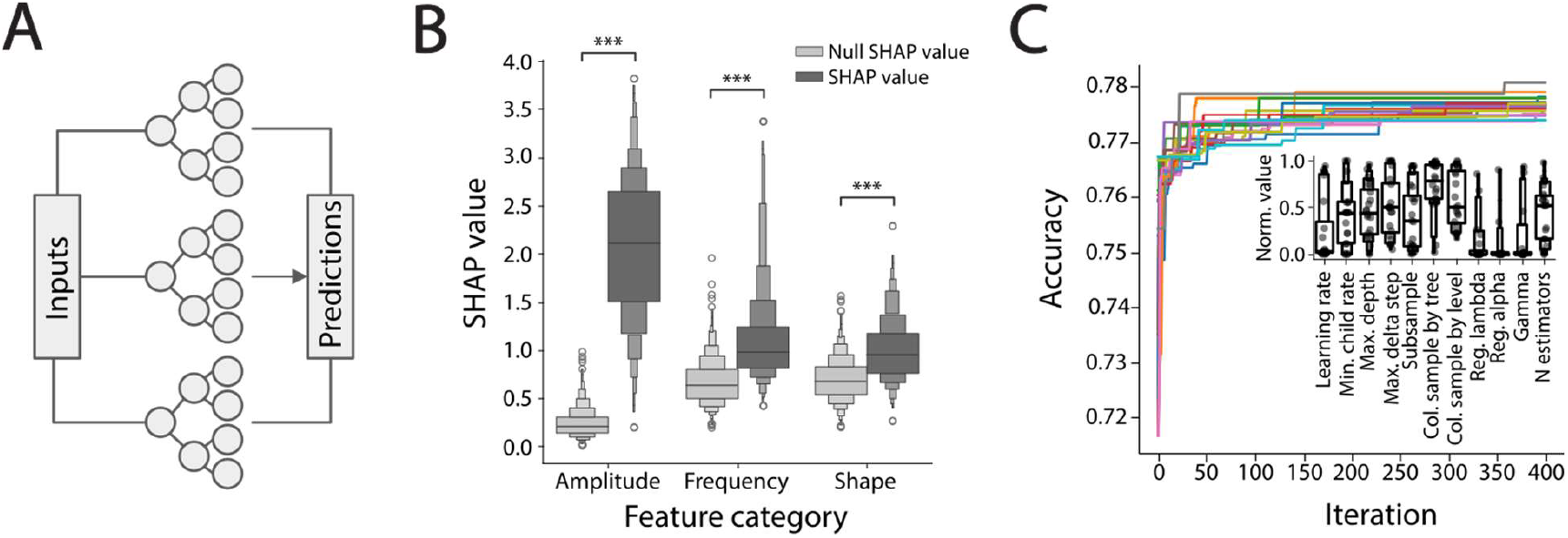
The XGBoost model accurately classifies EMG data. **A)** The XGBoost architecture represented as a set of tree-based models. **B)** Interrogation of feature importance using SHAP values shows that all sets of features show stronger contribution to predictions than expected by chance (Null SHAP values; ***: p < 0.001, Wilcoxon signed-rank). **C)** Classifier model performance is robust to hyperparameter specification—models trained without hyperparameter optimization exhibit high cross-validation accuracy and hyperparameter optimization results in only small accuracy gains. **Inset)** Distributions of final sets of hyperparameters (normalized to search range) following optimization. Parameters span almost the entire search range but some show tighter clustering.

This analysis confirmed that each of the eight features contributed significantly compared to control SHAP values generated by models trained on shuffled labels (p < 0.001 for all features, Wilcoxon Signed-Rank Tests). Since SHAP values are additive, we obtained a better understanding of the importance of each feature group by visualizing the summed values for each group of features. We found amplitude to be the feature with the highest absolute SHAP value, followed by frequency- and shape-related features. (**Fig. 2B**), but the differences between the importance of features within each group were not significant (i.e. all frequency features had equal importance to each other, as did all shape features, p > 0.1 for features in frequency group, and in shape group, ANOVA).

To assess the robustness of the model against specific hyperparameters (parameters determining model structure and training that do not get optimized as part of training); we tested whether the model’s performance was dependent on those settings by performing 20 repeats of hyperparameter optimization, randomly selecting starting hyperparameters anew with each repeat (in **Fig. 2C**, each repeat is represented by a different color). We found that the leave-one-animal-out cross-validation accuracy of the model, which is high even with the initial random setting (accuracy = 0.74±0.02, mean±sd; chance accuracy = 0.33), increases only slightly following hyperparameter optimization (~4% increase, 0.78±0.002, p < 0.001, paired t-test versus initial values). The distribution of hyperparameters from the 20 runs covered almost the entire search range for each hyperparameter, but nonetheless the model consistently converged to similar values of cross-validation accuracy (see **Table 2** for search space specifications, fraction of search space covered by final distributions = 0.93±0.05, mean±sd across hyperparameters)—that is, hyperparameter sets with widely varying values gave similarly high accuracies (although some hyperparameters did show tighter clustering, **Fig. 2C Inset**). Overall, this result demonstrates that the predictive capacity of the model reflects robust interpretability of the EMG data, rather than some idiosyncrasy of model definition and training.

We next contrasted the performance of the XGB classifier with that of BSA and QDA. **Figure 3A** shows, for all three classifiers, a representative example of predicted behavioral labels based on EMG waveforms, for trials of four tastes. Even casual inspection of the panels demonstrates that gapes are, as expected, consistently identified by all classifiers as occurring largely in response to the unpalatable tastant, quinine, and to appear, at earliest, 500ms following stimulus delivery. We compared the accuracy of each of the different algorithms’ classification of user-scored (“ground-truth”) gapes, and found that our XGB classifier predicted gape responses marginally better than QDA and significantly better than BSA (**Fig. 3B**; top left value in each matrix); despite the latter algorithms being specifically designed for that task (Li et al., 2016; Mukherjee et al., 2019; Sadacca et al., 2016), XGB performed better.

**Figure 3.**
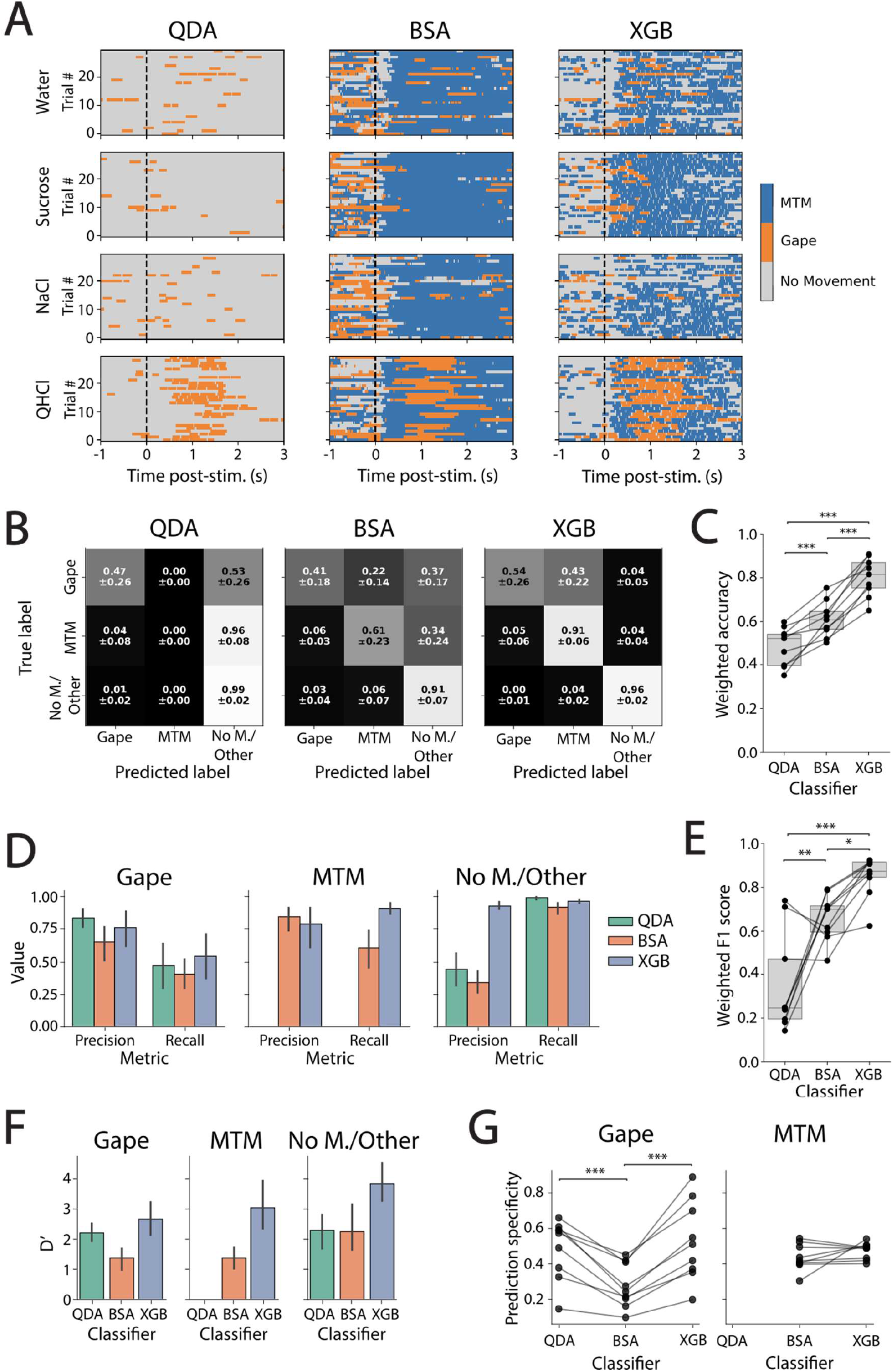
XGB outperforms previously used classifiers. **A)** Example predictions by QDA, BSA, and XGB classifiers for four tastants. **B)** Session-averaged confusion matrices of class-specific predictions for all three models (mean±STD). **C)** Weighted accuracy for all three models. XGB outperforms both QDA and BSA (Tukey post-hoc). **D)** Prediction precision and recall for all three models across label class. XGB performs on par with or better than QDA and BSA for all classes on both metrics. **E)** Weighted F1-score for all 3 models. XGB outperforms both QDA and BSA (Tukey post-hoc). **F)** D’ values for binary classifications. XGB consistently performs best for all class labels. **G)** Comparison of temporal specificity of rejection/ingestion-related responses (paired t-tests). * p < 0.05, ** p < 0.01, *** p < 0.001, for all panels

The remainder of the **Figure 3B** matrix shows how well the classifiers predicted “ground-truth” for other behaviors. Note that QDA is incapable of making MTM predictions, and that BSA generates “no movement” predictions by assuming that waveform frequencies <4 Hz are meaningless. **Figure 3C** summarizes these data, showing weighted accuracy (normalized for the very different proportions for behavioral classes in the dataset—i.e., the fact that the majority of identified behaviors were MTMs). Unsurprisingly, QDA performs most poorly, as it cannot account for an entire class (MTMs). Meanwhile, XGB performs best.

To compensate for the overall performance disadvantage that QDA’s missing MTM label places it at in relation to BSA and XGB, we compared all 3 algorithms in a binary classification task, using only the categories QDA is capable of predicting (i.e., gapes and “no movements”). All algorithms performed similarly well for binary classification of gapes, but for binary classification of “no movements,” we find XGB once again shows the best performance, followed by BSA, with QDA performing most poorly (see **Supplementary Figure 2**, also see below for D’ results on binary classification). Of course, neither this nor the above measures provide a full evaluation of classifier performance, because a truly competent model needs to have both high precision (the fraction of predicted hits that are “true” hits) and recall (the fraction of “true” hits that are not “missed”). An analysis of these two metrics (**Figure 3D)** shows XGB to perform on par with QDA with regard to gapes (both perform slightly better than BSA). Meanwhile, both BSA and XGB show high precision with regard to MTMs, but XGB performs far better with regard to recall (i.e., missing few true positives). Finally, for “no movements,” all classifiers have similar recall but only XGB shows high precision.

Given that high recall is of limited use without (at least reasonably) high precision (e.g., a classifier that gives all datapoints the same label will have perfect recall for that class but poor precision), the most rigorous evaluation involves aggregating the two measures–one such aggregation is a weighted F1-score. As shown in **Figure 3E**, QDA performs worst on this measure, BSA does slightly better, and the XGB classifier produces the highest F1-scores. Overall, XGB best balances recall with precision to identify known behaviors with significantly improved accuracy, outperforming both other algorithms. As a convergent test, we compared the classifiers using d’ (Fisher Discrimination Index, see Methods), which provides a bias-free measure of classification: derived from signal detection theory for binary classification tasks, d’ is calculated by Z-transforming the difference between true positive rate and the false positive rate; values between 1 and 2 are interpreted as evidence for weak performance while those higher than 2 indicate strong performance (Macmillan & Creelman, 2004). By this rubric, both QDA and XGB perform well on gapes, while BSA performs weakly. BSA also performs weakly on MTMs (due to poor recall) while XGB performs strongly. Finally, both QDA and BSA perform strongly on “no movements,” but XGB significantly outperforms both (**Fig. 3F**). In summary, XGB is the only algorithm for which d’ > 2 for all classes.

We performed one additional independent test of the prediction quality of the XGB classifier, assessing the fraction of taste-responsive movements (gapes and MTMs) that occurred post-stimulus delivery. Our reasoning was that rats should make almost exclusively “no movements” in the absence of a stimulus—gapes and MTMs should largely be confined to the post-stimulus time-period. Our test of this hypothesis (**Figure 3G**) shows that the XGB classifier produces significantly higher “post-taste specificity” than BSA for the prediction of gapes, performing on par with QDA (QDA vs BSA t(8) = 6, p < 0.001, XGB vs BSA t(8) = 6.6, p < 0.001, QDA vs XGB t(8) = 1.2, p =.277, paired t-tests for all comparisons), while XGB matched BSA for specificity of MTMs (t(8) = 1.3, p = 0.228, paired t-test; again, QDA is unable to predict MTMs altogether). In summary, only XGB performs well across the entire range of behaviors.

This validation, in conjunction with the fact that the XGB classifier also (verifiably) classifies categories of behaviors that previous algorithms cannot handle, gave us the confidence to apply XGB to 22 additional test sessions with AD EMG recordings (n = 9 animals; see Methods), and to use model predictions to characterize the structural and temporal properties of MTMs.

### MTMs are comprised of three distinct behavioral subtypes

The central goal of the current study is to explore whether MTMs are complex and non-uniform, and to test the hypothesis that the decision to consume is characterized by a change in MTM frequency. Since the MTM class as defined here is an amalgam of two behaviors (see Methods), it is reasonable to expect that they will not manifest as a single, uniform type. Although visual inspection alone (of video and EMG data) fails to reveal any such subtypes, we hypothesized that further analysis of the EMG features of XGB-classified MTMs would reveal distinct and consistent subclasses/clusters.

In preparation for these analyses, we linearly reduced the dimensionality of the feature data using Principal Component Analysis (PCA), a step that has been shown to improve model fitting quality (Aggarwal et al., 2014) while preserving most of the data’s original structure. All clustering analyses were thereafter conducted in 6-dimensional PCA space (reduced from 8 dimensions, retaining 90% of the variance; note, however, that the results below are unchanged by the use of the full 8 dimensional dataset). UMAP (a stochastic, non-linear transform that distorts distances) was used purely for visualization since it represents the data qualitatively.

To determine the best number of clusters to describe the MTM data, we applied Gaussian Mixture Modeling (GMM) with 1-14 clusters to the PCA-reduced waveform features in each individual test session. The number of clusters that best describe the data was identified using the Bayesian Information Criterion (BIC), a goodness-of-fit metric that accounts for model complexity to prevent overfitting. **Figure 4A** shows a representative example of this analysis, which reveals that MTM features, shown in UMAP space, are best described as consisting of three distinct clusters of behavior subtypes—note the sharp elbow in the BIC plot (**Fig. 4A inset**).

**Figure 4.**
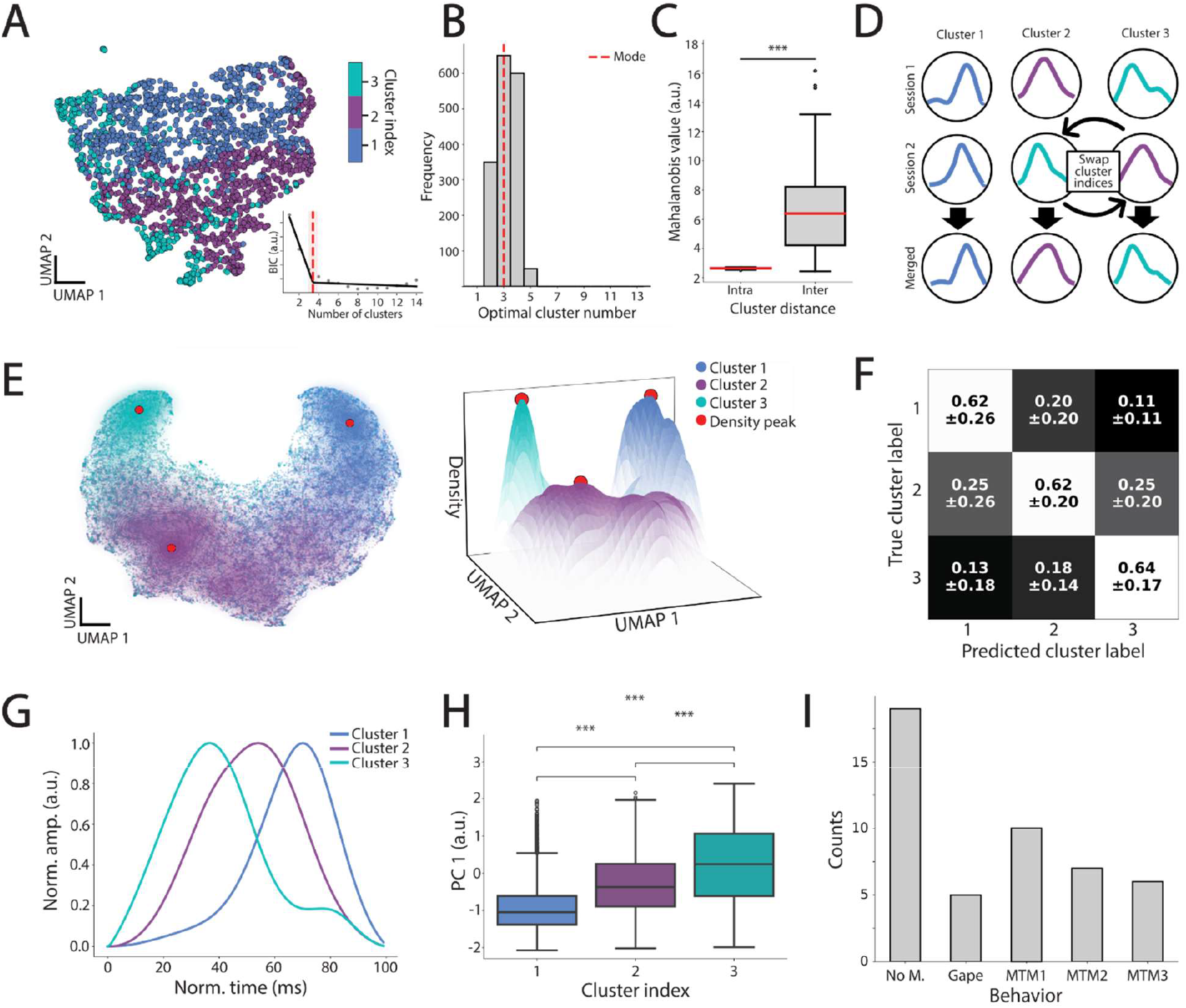
MTMs are comprised of three distinct behavioral subtypes. **A)** Example MTM waveforms in UMAP space, showing individual waveforms falling into 3 clusters. **Inset)** BIC values for 1-14 clusters indicating that three clusters are optimal for this dataset, based on the elbow method. **B)** Distribution of optimal cluster number across 50 iterations for each test session (mode = 3). **C)** Comparison of intra-cluster versus inter-cluster Mahalanobis distances indicates high separability of clusters for each session (*** p < 0.001, Kolmogorov-Smirnov test). **D)** Schematic depicting cluster index alignment across test sessions using cosine similarity. **E)** MTM waveforms across all test sessions projected into UMAP space, colored by cluster index label. (Left) Two-dimensional view (“the croissant”) highlighting the spread of the MTM distribution. (Right) Surface plot showing cluster densities with density peaks (red dots). **F)** Confusion matrix from a SVM (support vector machine) classifier showing robust cluster prediction accuracy for leave-one-animal-out cross-validation. **G)** Representative waveforms for each cluster, selected as those closest to the density peak. **H)** The first principal component (PC 1) values, which are derived from waveform shape, are significantly different across clusters (*** p < 0.001, Mann-Whitney U-Test). **I)** Counts of video-annotated lateral tongue movements classified into each behavioral category.

This conclusion was robust across multiple fits to the same dataset (50 repeats for each test session), multiple test sessions, and multiple rats (**Fig. 4B**; mean optimal number of clusters = 3.3). We therefore simplified subsequent analyses by restricting models to three clusters for every session. We added a validation step to confirm that this model restriction was reasonable, testing whether the clusters were truly distinct within sessions by comparing pairwise Mahalanobis distances between and within clusters; this test confirmed that inter-cluster distances were significantly larger than intra-cluster distances (D statistic = 0.94, p < 0.001, Kolmogorov-Smirnov test; **Fig. 4C**). Together, these analyses suggest that MTMs are a category umbrella over three subtypes of behavior (rather than being a simple amalgam of tongue protrusions and mouth movements).

To aligned cluster indices across sessions (which are randomly assigned during the initial clustering process, see Stephens, 2000 regarding “label-switching” in mixture models), we used a “greedy approach” (Cormen et al., 2001) that maximized the summed cosine similarity across all clusters from two test sessions, (see Methods and **Fig. 4D**) and continuing to merge sessions until all sessions were aligned. We visualized waveforms across all sessions by projecting each into UMAP space (**Fig. 4E**). This representation makes it clear that MTM waveforms across our entire dataset occupy overlapping yet distinctive regions with well separated density peaks (noted with red dots). To quantitatively check the distinctiveness and stereotypy of clusters across sessions, we trained a support vector machine classifier to predict cluster labels and tested it using a leave-one-animal-out cross-validation scheme. This classifier predicted cluster index significantly above chance (*t*(32) = 10.36, p < 0.001, one sample t-test; **Fig. 4F**), confirming that the three clusters are consistently distinct across animals. We refer to MTM clusters 1,2,3 as MTM1, MTM2, and MTM3 for the remainder of the manuscript.

This distinctiveness of clusters implies that waveforms in one cluster differ significantly from those in the others, and that this set of three waveform shapes are consistent from rat to rat. To directly test this implication, we first visualized the archetypal shape of waveforms in each cluster by plotting the waveforms closest to each cluster’s peak density. When viewed side by side, these waveforms exhibit clear differences in shape and appear to parametrically vary with regard to symmetry (**Fig. 4G**). To quantitively assess the apparent differences between the clusters, we performed Kruskal–Wallis tests, which revealed significant differences between clusters for all features (all p < 0.001). Note, however, that effect sizes varied from barely noticeable for most features (ε^2^ < 0.04), to moderate for Principal Component (PC) 2 and time to previous peak (ε^2^ =0.1 and 0.04, respectively), to relatively strong for PC 1 (ε^2^ = 0.23; see **Table 3**).

**Table 3.** Summary of Kruskal-Wallis test results for feature differences between clusters.

| Feature | H | p | $\epsilon^2$ | Effect size |
| --- | --- | --- | --- | --- |
| Duration | 1339.24 | < 0.001 | 0.0325 | Weak |
| Distance from last peak | 1842.74 | < 0.001 | 0.0448 | Moderate |
| Distance from next peak | 629.23 | < 0.001 | 0.0153 | Weak |
| Maximum frequency | 1112.94 | < 0.001 | 0.0270 | Weak |
| Normalized amplitude | 43.48 | < 0.001 | 0.0010 | Negligible |
| Principle component 1 | 9587.23 | < 0.001 | 0.2333 | Relatively strong |
| Principle component 2 | 4132.21 | < 0.001 | 0.1005 | Moderate |
| Principle component 3 | 194.32 | < 0.001 | 0.0047 | Negligible |
| Amplitude | 29.67 | < 0.001 | 0.0006 | Negligible |
| Area under the curve | 8.71 | < 0.001 | 0.0001 | Negligible |
| Full width half max | 206.71 | < 0.001 | 0.0049 | Negligible |
| Rising slope | 281.80 | < 0.001 | 0.0068 | Negligible |
| Falling slope | 552.79 | < 0.001 | 0.0134 | Weak |
| Symmetry | 4804.77 | < 0.001 | 0.1169 | Moderate |
| Skew | 1464.57 | < 0.001 | 0.0356 | Weak |
| Bimodality | 1754.77 | < 0.001 | 0.0427 | Moderate |

As PC1 had the greatest effect size, was cluster-specific (p < 0.001 for all pairwise comparisons, Mann-Whitney U-Test; **Fig. 4H**), and is derived directly from waveform shape, we dissected how the shapes differed by measuring an additional set of descriptive features derived from each waveform’s shape: area under the curve, rising slope, falling slope, full width half max, symmetry, skewness, and bimodality. Kruskal–Wallis tests revealed significant differences between clusters for all metrics (p < 0.001 for all comparisons), with symmetry and bimodality having the largest effect sizes (ε^2^ = 0.12 and ε^2^ = 0.04, respectively; see **Table 3**). This quantification confirms the differences in symmetry visually apparent in the representative waveforms in **Figure 4G** and suggests that MTM1 and MTM3 are not simply mirror images. To further validate this, we mirrored the waveforms of MTM1 and re-trained a support vector machine on the derived principal components. The classifier was able to accurately predict the clusters labels in a leave-one-animal-out cross-validation scheme (mean accuracy = 0.7), performing significantly above chance (t(32) = 20.03, p < 0.001, one-sample t-test), indicating that the 3 waveform shapes are truly distinct from one another, irrespective of skew direction.

Each AD contraction is part of the mechanism underlying a single orofacial movement, working in concert with other lingual and masticatory muscles. Given the presence of three distinct clusters of AD waveforms within MTMs, we conclude that each MTM is a distinct behavioral subtype which is found consistently across sessions and rats. Although we cannot infer directly how waveform cluster differences in symmetry and bimodality reflect visual differences in orofacial movements (again, these MTMs were difficult to distinguish by eye), the results suggest meaningful differences in the temporal dynamics of the muscle contractions—differences that have been shown using biophysical modelling to impact the shape of the movement, particularly in the context of other oromotor muscles’ activity (see Discussion, and Ross et al., 2024). In this framework, we considered the possibility that the three clusters cleanly reflected the three ingestive behaviors originally proposed by Grill and Norgren (1974a): mouth movements, tongue protrusions, and lateral tongue movements. More specifically, we asked whether one of the three clusters might reflect lateral tongue movements by testing whether the ~5% of behaviors identified as lateral tongue movements by the video coders (**Fig. 1F**), which were too few in number to train as a distinct behavioral class for the classifier, would be assigned to a particular MTM cluster.

The results of this analysis (**Fig. 4I**) disproved this hypothesis: despite being associated with sizeable AD activity, lateral tongue movements were assigned to the “no movement” category 2-3 times as often as they were to any MTM category. Furthermore, the likelihood that a lateral tongue movement would be sorted into one of the MTM clusters was similar to its likelihood of being categorized as a gape. Thus, we conclude that XGB allows us to identify a novel trio of MTM subtypes, each of which involves a distinctive time course of AD activation.

### Frequency of MTM subtypes changes between the beginnings and ends of trials

If the decision to ingest a taste primarily involves organizing MTMs to facilitate swallowing, our central prediction is that the frequency of at least one MTM subtype should change while the rat is preparing to ingest. We examine three competing possibilities: 1) a specific subtype of MTMs is associated with the onset of ingestion (analogous to the sudden appearance of the single behavior—gaping—for rejection decisions; see Figure 3A); 2) there is a broader behavioral pattern shift underlying ingestion which is characterized by frequency changes of multiple MTM subtypes; or 3) the null hypothesis that MTMs do not change in a manner suggestive of involvement in the consumption decision.

We first tested for any broad changes in MTMs across time by separating trials into halves—0-800 ms and 801-1600 ms post-taste—a dividing line chosen based on previous work on the timing of rejection-related behaviors (i.e., gaping; Travers & Norgren, 1986; Sadacca et al., 2016; Mukherjee et al, 2019). For each test session, we then reduced the dimensionality of MTM features in each part of the trial and visualized the results using UMAP projection. This presentation revealed distinct differences between the “first” and “second trial-half” distributions of MTM features (**Fig. 5A**). We evaluated the significance of these differences by comparing the difference in the first-and second-half distributions using a histogram-difference metric, repeating the processing 10 times to ensure robustness to stochasticity (see Methods). The median p-value for all sessions was below 0.05, and in only 6 of 33 sessions were there outliers above that criterion (**Fig. 5B**), confirming that overall the MTMs produced before the approximate time of palatability-related firing are distinguishable from those produced after.

**Figure 5.**
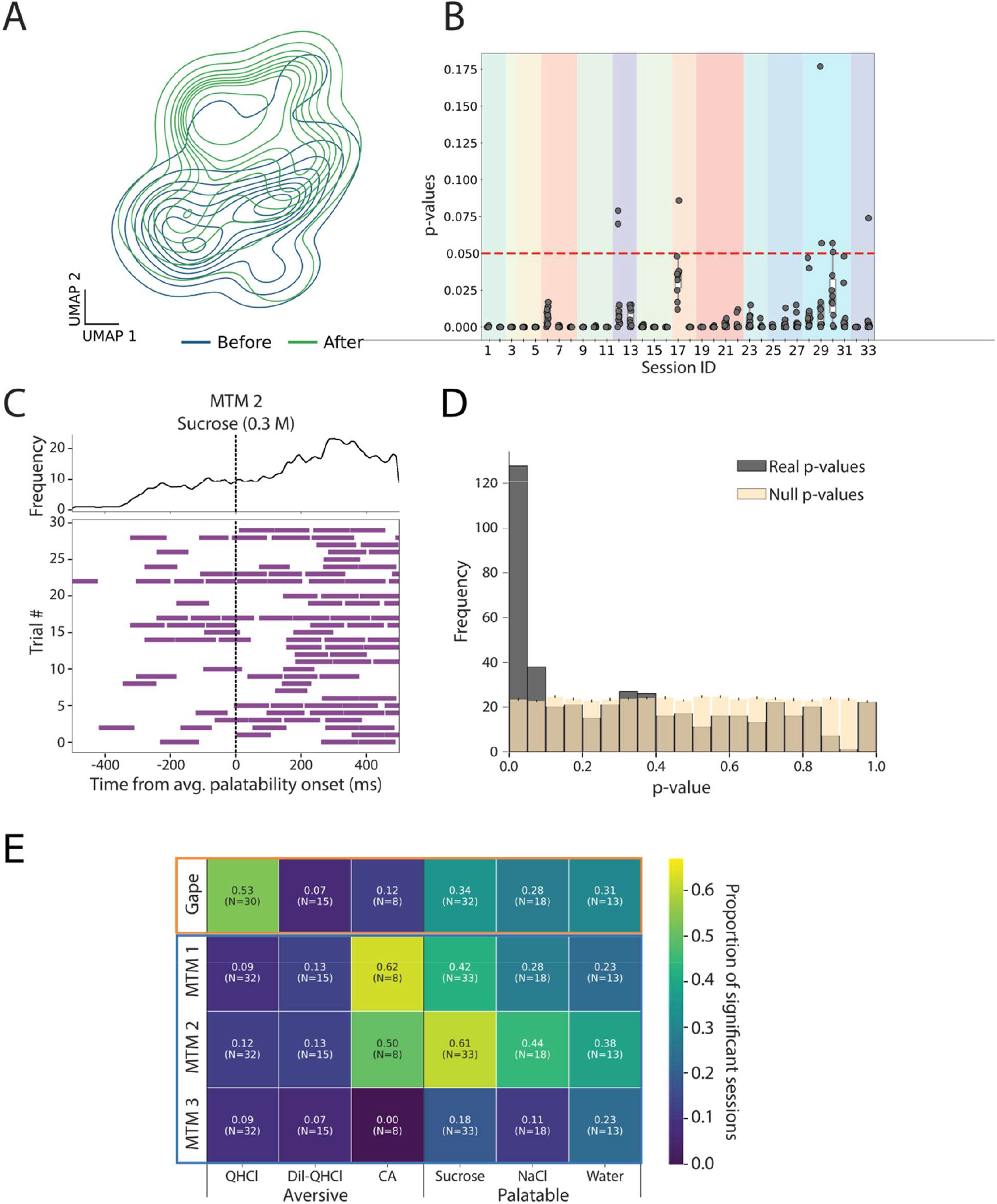
MTM subtype frequencies shift across the taste response. **A)** Example contour plot of UMAP-reduced MTM feature distributions categorized by their timing relative to average palatability activity onset (“before” [blue] versus “after” [green] 800 ms post-stimulus delivery).**B)** Session-wise p-values quantifying the shift in MTM feature distributions before versus after average palatability activity, calculated using bootstrap resampling. The median p-value across multiple repeats for all sessions is <0.05. Colors denote sessions from the same animal. **C)** <u>Bottom:</u> Raster plot, and <u>Top:</u> mean rate of occurrence of MTM cluster #2 across 30 trials of sucrose in an example test session, aligned to the average palatability activity onset (800 ms post-stimulus delivery). **D)** Distribution of Poisson means test p-values across all tastant-behavior comparisons for every test session (gray) compared to a null distribution (yellow). **E)** Tastant-behavior matrix, with brightness of color indicating the proportion of significant test sessions showing broad changes in behavior frequency.

This result could be observed in the production of individual behaviors. **Fig. 5C** illustrates, for example, the times at which one rat produced MTM cluster #2 (MTM2) across 30 sucrose trials in a single session. While the waveform could be found at almost any post-stimulus latency, the occurrence of MTM2 increased dramatically after the 800-ms dividing line. This example was representative: the significance of behavior changes within trials (Poisson means tests for 473 comparisons across all sessions, tastants, and behaviors) was highly skewed toward low p-values (**Fig. 5D**), with the fraction of p-values<0.05 significantly exceeding the proportion expected by chance (p < 0.001, binomial test).

.Having confirmed the presence of significant changes in the frequencies of behavior across time, we moved on to testing how these changes are distributed across tastes and behaviors. Rejection decisions have repeatedly been shown to involve the sudden appearance of gaping alone (e.g., Grill & Norgren, 1978a; Sadacca et al., 2016). We hypothesized that within-trial shifts in response to aversive tastants would therefore be driven primarily by gapes, whereas palatable tastants would show mid-trial changes in multiple MTMs. We further predicted that the consistency of the pattern change should scale with the tastant’s valence; that is, strongly aversive or palatable tastants will elicit more robust changes in oromotor behaviors.

A tastant-by-behavior significance matrix, showing the fraction of significant test sessions for each tastant-behavior combination (**Fig. 5E**), confirmed these hypotheses. All tastant-behavior combinations were significant above chance (that is a fraction >0.05 of tests were significant, with a single exception of MTM3 to citric acid), highlighting the robustness of the change. Gapes proved to be the behavior that changed most consistently in response to highly aversive quinine (QHCl), recapitulating previous findings (Mukherjee et al., 2019; Sadacca et al., 2016). Dilute quinine (Dil-QHCl), expectedly, showed weakly consistent changes across all behaviors (Grill & Norgren, 1978a). In addition, aversive citric acid unexpectedly elicited a highly consistent change to MTM1 frequency, a result we address in the Discussion.

For palatable tastants, meanwhile, we observed highly consistent changes across all three MTM clusters. For MTM1 and MTM2, the consistency of these changes scaled with palatability: sucrose elicited the most consistent changes (both MTM1 and MTM2 showed high fractions of significant comparisons, 0.42 and 0.61 respectively); within-trial MTM changes to slightly less-palatable sodium chloride [NaCl] were somewhat less robust, as were those in response to water (which only showed relatively high fractions from MTMs, 0.44 and 0.38 for each taste respectively). Altogether these results confirm that taste processing culminates in reliable but broad changes in behavior as part of ingestion, and extend these conclusions—beyond the previously demonstrated onset of gaping to quinine, there is reorganization in the numbers of all three types of MTMs as rats prepare to ingest palatable tastants.

### The frequency of MTMs specifically changes just following the Gustatory Cortex transition to palatability-related firing

The above results motivate and support our most risky hypothesis, in that they reveal broad changes in the occurrence probability of MTM clusters elicited by palatable tastants as rats approach the decision to consume. We propose that these changes in fact represent the reaching of that decision, and more specifically that they occur just after the time at which GC ensemble taste responses transition into a period of palatability-related firing (Katz et al., 2001; Sadacca et al., 2012), as has previously been found to signify the rejection decision made in response to aversive tastes (Sadacca et al., 2016; Mukherjee et al., 2019).

To rigorously test this hypothesis in a manner that does not presuppose a specific time point for a behavioral change—an important consideration, given the wide trial-to-trial variability in both decision and neural transition latencies (Sadacca et al., 2016; Mukherjee et al., 2019)—we analyzed sessions in which EMG and GC single-neuron taste responses were simultaneously recorded (22 test sessions across 9 animals), fitting changepoint models to time series of behaviors to determine when transitions in the occurrence probabilities of the behavior occur (allowing us to evaluate changes in all MTM subtypes collectively) and comparing the timing of these inferred changes to onsets of palatability-related ensemble activity in GC (in the same trial).

Performing this analysis required that we first determine the number of states that best describe the dynamics of the classified orofacial behaviors. We fit change point models with different numbers (2-8) of states to 2000 ms of post-stimulus activity and compared the models using the ELBO (**E**vidence **L**ower **Bo**und, not to be confused with the “elbow method” used to evaluate BIC in the previous section), the metric used to fit Bayesian models using variational inference (see Methods). Examined in this way, models with 3 states (that is, 2 changepoints) provided by far the best fit to the data (**Fig. 6A**; p < 0.005 for all pairwise comparisons, t-test with Bonferroni correction); hence the 3-state model was used for all subsequent analyses. **Fig. 6B** shows a single-session representative example of behaviors observed across all tastes (**Fig. 6Bi**) along with the timing of the states and changepoints inferred by the model (**Fig. 6Bii**). Note that the timing of the changepoints varies from trial to trial, even within a taste, as has been shown for the GC ensemble transition into palatability/decision coding (Sadacca et al., 2016). The probabilities of each behavior type occurring in a particular state (at a particular moment in time) for any given taste (**Fig. 6Biii**) further demonstrate that the composition of states shifts over time, consistent with broader changes in movement patterns.

**Figure 6.**
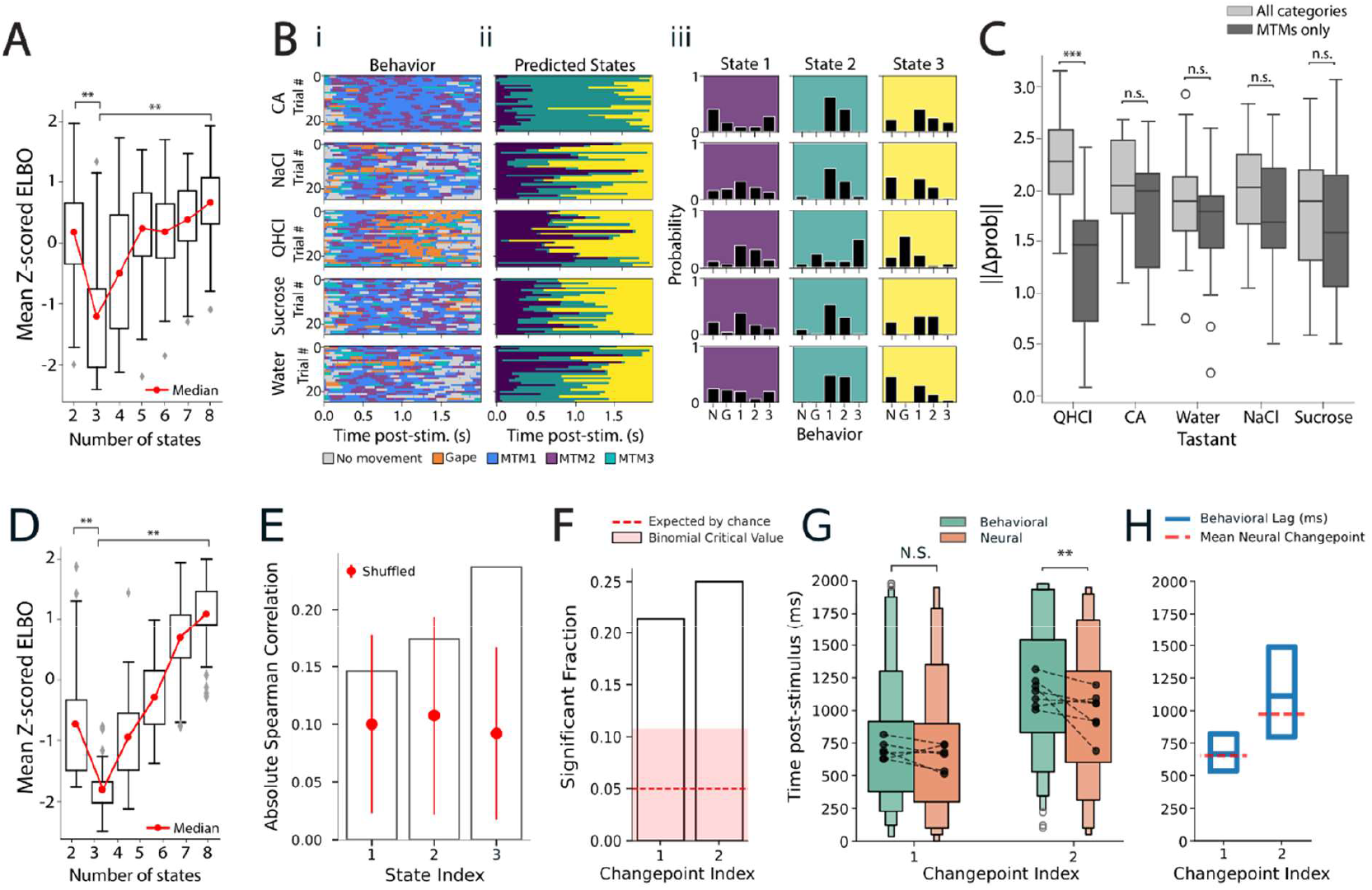
Emergence of palatability encoding in GC neural activity precedes changepoints in MTM occurrence. **A)** Comparison of changepoint models with 2-8 states applied to timeseries of oral behaviors reveals that 3-state models best describe behavior dynamics in 2000 ms post-stimulus delivery (** p < 0.01, pairwise paired t-test with Bonferroni correction). **B)** Representative example of inference performed by changepoint model. **i)** Raster plot of behaviors across tastes; **ii)** State sequences inferred by changepoint model; **iii)** Histograms of probabilities for each behavior per state and taste inferred by changepoint model. N = no movement, G = gape, 1, 2, 3 = MTM cluster 1, 2, and 3, respectively. **C)** Comparison of normalized magnitudes of difference between inferred behavior probability vectors for subsequent states, including either all categories or MTMs only. Only quinine (QHCl) is affected when gapes and no movements are removed (*** p < 0.001, t-test with Bonferroni correction). **D)** Comparison of changepoint models with 2-8 states applied to timeseries of electrophysiological responses; 3-state models best describe neural dynamics in 2000ms post-stimulus delivery (** p < 0.01, pairwise paired t-test with Bonferroni correction). **E**) Only activity in the 3^rd^ neural state (i.e., the state that follows the 2^nd^ changepoint) is significantly correlated with palatability. **F)** The neural and behavioral latencies of both changepoints 1 & 2 are significantly correlated in a higher number of sessions than expected by chance **G**) Comparison of lags between neural and behavioral changepoints latencies shows a significant neural-to-behavioral lag for the 2^nd^ changepoint but not the 1^st^. Black dots and connecting lines show session-average latencies. **H)** Same data as in G) plotted as lags of behavioral changepoints against mean latency of neural changepoint to highlight difference of lags between both changepoints.

Furthermore, a 2-way ANOVA (states x tastes) on ELBO values for all sessions showed significant differences between the number of states (F(6)=172.9, p < 0.001) but not between tastes (F(4)=0.005, p = 0.99), and we found no significant differences between tastes for the ELBO in 3-state models (p > 0.1 for all pairwise comparisons between tastes, t-test with Bonferroni correction). These results make it clear that the 3-state solution describes the behavioral results of taste processing in general (rather than being the response to a particular taste or subset of tastes).

As convergent tests of our hypothesis that behavior change points in responses to palatable tastes are driven primarily by MTMs (and not gapes), we performed a pair of analyses. First, we confirmed that the probabilities of MTM clusters change with state (3-way ANOVA [state, taste, MTM cluster]: state: F(2)=1.21, p < 0.001, taste: F(4)=0.28, p<0.01, MTM cluster: F(2)=3.31,p<0.001), in line with results in the previous section (see **Supplementary Figure 3** for a full breakdown of each behavior across states and tastes). To complement this test, we instead assessed the extent to which removing gapes and “no movements” impacted the difference between sequentially-appearing states. Logically, if differences between states for palatable tastants are driven largely by gapes and “no movements” rather than MTMs, removing gapes and “no movements” should make the structure of states for palatable tastants more similar, As expected, the results of this analysis were taste-specific (2-way ANOVA; category [all vs. MTM only]: F(1)=0.15, p = 0.52, category*taste: F(4)=10.4, p < 0.001): **Figure 6C** shows that the difference in the structure of subsequent states was reduced when gapes were removed only for quinine (p < 0.001, t-test with Bonferroni correction); meanwhile, gape and “no movement” removal had no impact on the magnitude of the difference between states for any other taste (p > 0.1, t-test with Bonferroni correction). The fact that gapes only strongly affect the decision-related behavior change in response to quinine means that MTMs (in a distributed manner across all subtypes) are the primary drivers of behavior transitions in response to palatable tastes.

Next, we brought analogous analyses to bear on neural ensemble data, showing, as expected (Sadacca et al., 2016; Mahmood et al., 2023; 2026), that neural responses were also best described using 3-state models (**Fig. 6D**, p < 0.005 for all pairwise comparisons, t-test with Bonferroni correction, mean latencies, changepoint 1: 653ms, changepoint 2: 973ms), and that the last of these states (the average onset of which occurred just less than 1.0s into the response; see below) uniquely carried palatability-related firing (**Fig. 6E**). These results further motivated and facilitated the test of our ultimate hypothesis, which is that the within-trial change in MTM production actuates the consumption decision reflected in the GC ensemble transition into palatability-related firing. This hypothesis comprises two specific predictions: 1) that the timing of the late change in MTM production should be correlated with the timing of the appearance of the palatability-related GC ensemble state in single trials; and 2) that the neural transition should lead the behavioral transition by a few hundred millseconds (see Sadacca et al., 2016).

A comparison of the latencies of the inferred changepoints in behavior and neural activity, accomplished using Pearson’s R for single recording sessions, supported our first prediction. This analysis (**Fig. 6F**) showed that, across the entire dataset, the number of significant correlations (alpha=0.05) in behavioral and neural transition times was higher than expected by chance according to a Binomial test (changepoint 1: p=0.002, changepoint 2: p<0.001), confirming that both transitions in behavioral and neural responses are strongly aligned in time in single trials. Analysis of lag structure (375 trials pooled across 9 sessions), meanwhile, revealed that the while there was no significant neural-behavioral lag in the 1^st^ changepoint (a result suggesting either common source or tight system coupling; see Discussion, mean lag = 50.7ms, t=1.466, p=0.144, paired t-test), the 2^nd^ neural transition reliably preceded the change in MTMs (**Fig. 6G**; mean lag = 199.6ms, t=4.816, p<0.001, paired t-test). The sudden onset of palatability-related firing in GC, which has been shown to predict and drive gaping to aversive tastes (Sadacca et al., 2016; Mukherjee et al., 2019), is appropriately timed to drive the changes in MTMs, thereby confirming that the trio of MTMs is involved in the decision to ingest.

We obtained qualitatively similar results when directly comparing average lag per-session (n=9 sessions, changepoint 1: p=0.127, changepoint 2: p=0.035, paired t-test, see paired black lines in **Fig. 6E**), and when we compared the number of single-session paired t-tests showing significant differences (changepoint 1: p=0.735, changepoint 2: p=0.031, 1-tailed Binomial test on count of significant single-session paired t-test). Hence, we conclude that the 2^nd^ changepoint uniquely demonstrated significant and reliable brain-to-behavior lag.

We conclude that this behavioral evidence of an ingestion decision, as detected by the changepoint model, is associated with the same neural transition in GC taste responses that has previously been shown to be responsible for the onset of gaping behavior, and that an ensemble of MTMs actuate the decision.

## Discussion

### Detection of orofacial movements using machine learning on EMG activity

Having a stimulus on the tongue necessitates a binary decision: to ingest or reject. The fact that the choice is binary does not however mean that it is simple. Variations in orofacial movements reflect subtleties of a rat’s palatability judgment—initially non-taste specific, then switching to aversive or hedonic—and underscore the necessity of understanding the full repertoire of movements to understand how the brain switches between these behavioral states. Grill and Norgren (1978a) proposed that the consummatory decision is actuated by highly stereotyped behaviors such as the gape, the primary rejective orofacial behavior, which is easily detected both visually and *via* EMG activity of the anterior digastric (AD; jaw opener) muscle. This distinctiveness has motivated multiple investigations into how the taste system drives gaping (Dinardo & Travers, 1994; DiNardo & Travers, 1997; Kinzeler & Travers, 2008; Li et al., 2016; Mukherjee et al., 2019; J. Travers et al., 2000). Meanwhile, similar studies probing the signals leading to the onset of the ingestive response are fewer, due in large part to the relative complexity of and lack of consensus regarding the two primary ingestive-related behaviors: mouth movements and tongue protrusions (Grill & Norgren, 1978a). In this study, we have leveraged recent advances in computational analysis in a reexamination of behavioral responses to palatable tastes, and in the process revealed novel insights into how the evolution of behavior within a tasting trial is linked to activity in Gustatory Cortex (GC).

To perform these analyses, we first trained a machine learning classifier to detect and classify individual orofacial movements from AD electromyographic (EMG) activity. We targeted the AD because it contracts as part of the execution of every movement cycle, with kinematics that show significant differences in duration between gaping and licking (J. B. Travers & Norgren, 1986)—of course, every movement involves AD acting in concert with other lingual and masticatory muscles, the activity of which necessarily influence one another—and because the muscle is readily accessible for EMG implantation. We chose to study movement using EMG activity not only because it allowed us to collect these data in freely moving rats but, more importantly, because it allowed us to avoid the inherent difficulties in distinguishing mouth openings from tongue protrusions in video recordings (in which the appearance of these behaviors is similar). With this in mind, we performed data labelling by defining an umbrella category of rapid mandible openings and closings with or without tongue extension—called mouth or tongue movements (MTMs)—that could then be parsed into finer subtypes (see below) based on features in the EMG signals.

The XGBoost classifier (XGB) presented in this paper outperformed previous algorithms on multiple metrics even when validated on detection of gapes, the behavior that these previously published classifiers were specifically designed to detect (Li et al., 2016; Mukherjee et al., 2019). This performance improvement almost certainly reflects XGB’s use of a suite of features—notably amplitude, frequency, and waveform shape—that together enhance discrimination of EMG waveforms generated by different behaviors. We also note that the XGB classifier performed ~1000x faster than BSA, and close to QDA speeds (**Supplementary Fig. 4**), making it suitable for processing larger datasets, and potentially even allowing studies utilizing online behavioral classification.

### Detection of three mouth or tongue movements (MTMs) subtypes

Our clustering analysis revealed that MTMs consistently group not into the expected two stereotyped clusters but rather into three novel clusters that are robust across sessions and animals. Each cluster represents unique temporal dynamics of the AD contraction, likely shaped by interactions with other orofacial muscles to produce three distinct movements which we termed MTM subtypes. Aa a group, they are distinct from the three ingestion-related behaviors originally described by Grill & Norgren (1978a). At this point in time, we cannot say what the differential function of each behavior is; we don’t know how they differ from one another with regard to moving fluid around the mouth, but we would speculate that together in a particular configuration (that is achieved with the mid-trial change in behavior) they function to facilitate preparation for swallowing the palatable fluid.

Investigating differences between clusters, we found that each cluster’s waveform shape was unique and stereotyped. While not perfectly distinct categories, they are well described as parts of a tri-modal distribution of movement patterns. This interpretation is consistent with the model proposed by Kaplan and Grill (1995) whereby mouth movements and tongue protrusions exist along a continuum of tongue extension.

Confirming how these clusters map onto specific mouth shapes remains challenging because all MTMs look the same in our video records and only become detectably different with the high temporal resolution of EMG. It is also possible that the speculative largest distinguishing feature, tongue movement, occurs primarily intraorally and thus was obstructed from view. Future work may begin to address this *via* video-based data mining in a head restrained paradigm (Bollu et al., 2020; Inaba et al., 2025) or biomechanical modelling of a broader EMG profile of more orofacial muscles. Regardless, our data support the hypothesis that these distinct AD contraction patterns give rise to movements that, across different coordination patterns, support both the non-taste-specific and ingestive-related phases of the taste response to differentially transport fluid around the oral cavity. (Ross et al., 2024).

Our data indicate that there is no single ingestion behavior directly analogous to gapes. Instead, ingestion involves coordination among a set of three MTM subtypes. We speculate that upon taste delivery, the rat may initiate a “sampling” phase characterized by one MTM pattern followed by an “ingestion” phase characterized by another (see Ross et al., 2024). It is only by considering the ingestive response as a pattern of multiple behaviors, through analyses such as changepoint modelling, that we were able to elucidate how the response evolves in relation to GC activity (see below). Future work could refine behavioral categorization by recording simultaneously from multiple muscles, with the extrinsic tongue muscles being an obvious first candidate (J. B. Travers & Norgren, 1986). Such an investigation would be able to definitively address whether the different subtypes of MTMs we observe form part of different sequences of muscle movements. The use of tools such as seqNMF (Mackevicius et al., 2019) could allow future investigations to detect sequential motifs in the behavior time series to characterize the patterns of MTM subtypes occurring within the states of the ingest response.

### The relevance of GC activity to ingestive decisions

Having established that there is an ensemble of three distinct MTM subtypes, we went on to show that there were broad significant changes in the ensemble response to all tastants between the early (0-800 ms) and late (801-1600 ms) post-delivery periods. In agreement with previously published findings (Li et al., 2016; Mukherjee et al., 2019; Sadacca et al., 2016), quinine trials were characterized by the consistent appearance of gapes across this boundary; meanwhile, palatable tastants elicited within-trial dynamics for all three MTM clusters, the consistency of which (particularly for MTM1 and MTM2) scaled with palatability. The fact that a strongly valenced tastant elicits a more reliable and robust behavioral response accords well with previous results (Grill & Norgren, 1978a; Spector et al., 1998; J. B. Travers & Norgren, 1986). Overall, these findings confirm that rejection and ingestion responses form a binary decision, and while nuanced behavioral patterns underlie each side of the decision based on the palatability of the tastant, the choice to ingest is marked by a shift in the behavioral pattern across the ensemble of MTMs.

Unexpectedly, citric acid also elicited uniquely consistent within-trial changes to MTM1 frequency at the time of consumption decisions. Although canonically considered to be aversive tastes, there are reports of acids eliciting mouth movements and/or tongue protrusions in rats and hamsters (Brining et al., 1991; Sasamoto et al., 2001; S. P. Travers, 2002). For reasons that remain unclear, sour tastants may elicit a rejection behavior pattern distinct from that elicited by quinine that consists of a mixture of ingestion and rejection-related behaviors; a detailed test of this hypothesis is needed, but is outside of the scope of the current study.

Finally, we directly tested whether the transition to palatability-related activity in GC modulated ingestion-related behaviors—previous work has established a causal relationship between this GC ensemble firing-rate transition and gape-driven rejection (Mukherjee et al., 2019; Sadacca et al., 2016), but it was unclear whether this result had specifically to do with aversion or represented an intrinsic facet of taste processing. Application of changepoint modelling to EMG activity revealed two time points in the first two seconds of the taste response at which there is a rapid shift in the behavior pattern of MTMs. The second transition matches the prior analyses, reflecting a change in behavior occurring soon after the middle of the trial. This latter transition proved coupled to the transition to palatability-related firing in GC (established in an entirely independent analysis), and followed that neural transition by approximately 200 msec. This result, which dovetails perfectly with previous findings regarding gaping (Mukherjee et al., 2019; Sadacca et al., 2016), suggests that GC—and in particular the sudden transition from identity-specific to palatability-related firing—is involved in both ingestion and rejection decisions. Whether or not ingestion and rejection decisions are transmitted to motor pattern generators by separate routes (Jin et al., 2021), GC is an integral part of both. Given that GC is a critical part of this decision and that the decision is ultimately actuated by a central pattern generator in the medullar reticular formation which has been shown to be responsible for both rejection and ingestive oromotor rhythmic patterns (Chen et al., 2001; DiNardo & Travers, 1997; Grill & Norgren, 1978b; Moriyama, 1987; Travers et al., 2000; Venugopal et al., 2007), the present results suggest that the two decisions are governed by the same underlying neural mechanisms.

