## Supplemental Figure 1 for "The Ingestive Response Reflects Neural Dynamics in Gustatory Cortex"

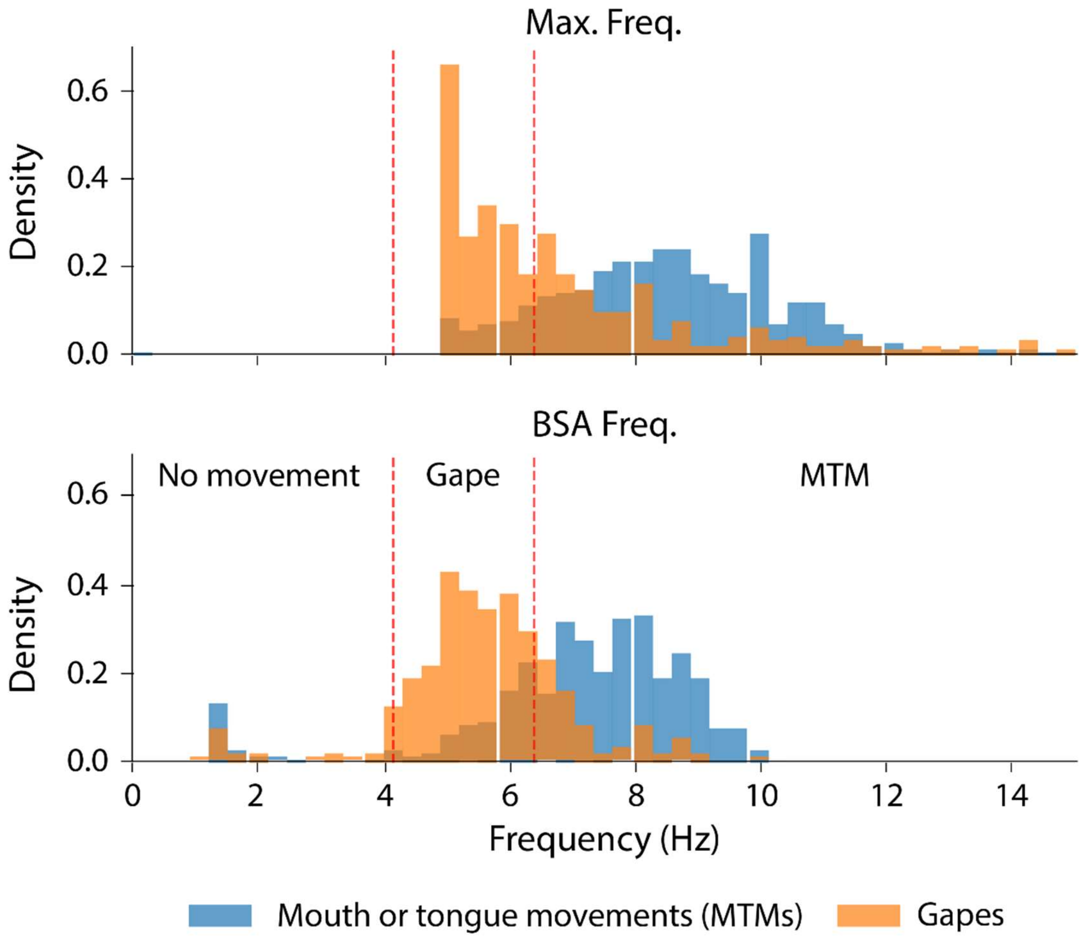


**Supplementary Figure 1**: Waveform frequency of scored gapes and MTMs accords well with thresholds set by previous work (Mukherjee et al. 2019), corroborating accuracy of scoring. Vertical red dotted lines indicate 4.15 and 6.4 Hz.
