## Supplemental Figure 2 for "The Ingestive Response Reflects Neural Dynamics in Gustatory Cortex"

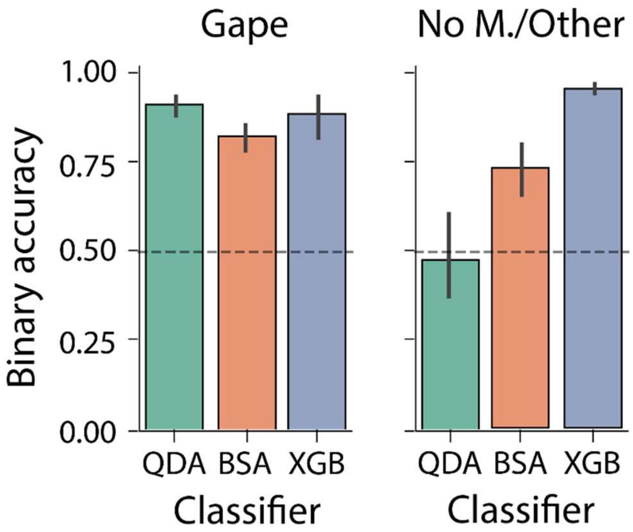


**Supplementary Figure 2**: XGB performs on par with, or outperforms QDA and BSA on binary classification for all 3 classes of oral behaviors.
