## Supplemental Figure 3 for "The Ingestive Response Reflects Neural Dynamics in Gustatory Cortex"

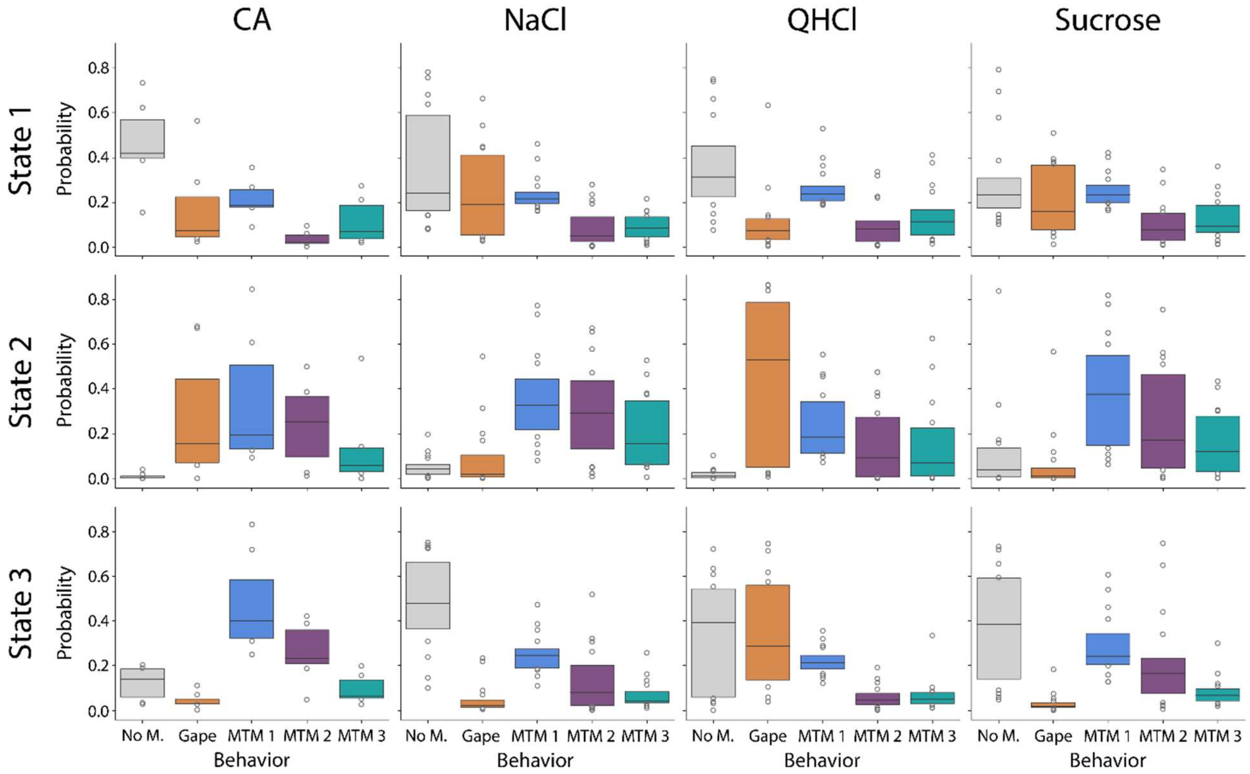


**Supplementary Figure 3**: Cross-session occurrence probability of all behaviors (and MTM subclasses) for inferred states across all tested tastants. Consistency of these probabilities across multiple sessions lends confidence to the robustness of state inference and the consistency of behaviors within tastes.
