## Supplemental Figure 4 for "The Ingestive Response Reflects Neural Dynamics in Gustatory Cortex"

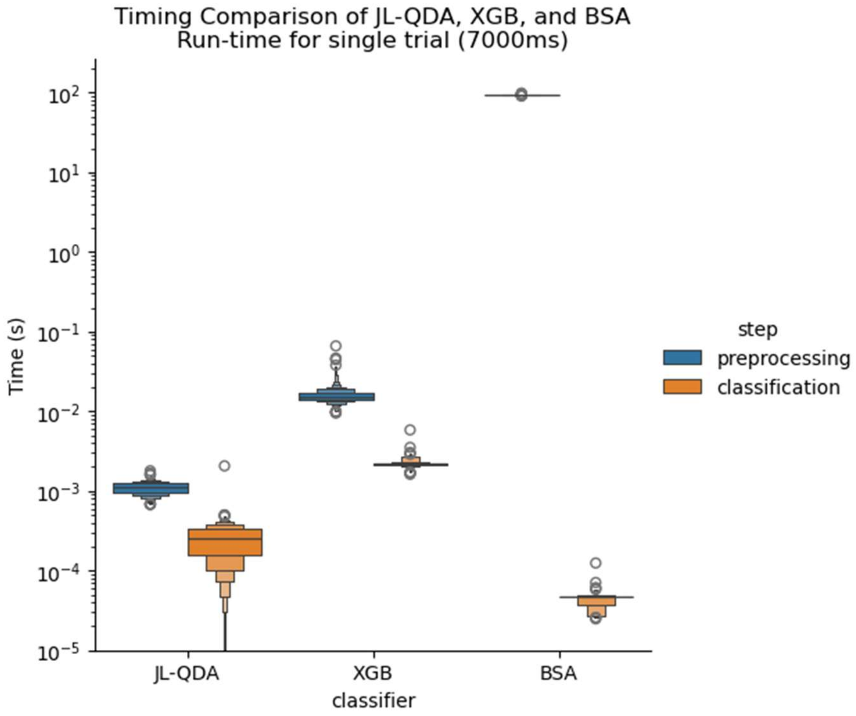


**Supplementary Figure 4:** XGB, despite being a more complex classifier, performs rapidly, processing single-trial (7000ms duration) data in <0.1sec. Both XGB and QDA perform >1000x faster than BSA.
